# Co-option of the Limb Patterning Program in Cephalopod Lens Development

**DOI:** 10.1101/2021.04.22.441006

**Authors:** Stephanie Neal, Kyle J. McCulloch, Francesca Napoli, Christina M. Daly, James H. Coleman, Kristen M. Koenig

## Abstract

Across the Metazoa, similar genetic programs are found in the development of analogous, independently evolved, morphological features. The functional significance of this reuse and the underlying mechanisms of co-option remain unclear. Here we identify the co-option of the canonical bilaterian limb pattering program redeployed during cephalopod lens development, a functionally unrelated structure. We show radial expression of transcription factors *SP6-9/sp1, Dlx/dll, Pbx/exd, Meis/hth*, and a *Prdl* homolog in the squid *Doryteuthis pealeii*, similar to expression required in *Drosophila* limb development. We assess the role of Wnt signaling in the cephalopod lens, a positive regulator in the developing limb, and find the regulatory relationship reversed, with ectopic Wnt signaling leading to lens loss. This regulatory divergence suggests that duplication of SP6-9 in cephalopods may mediate this co-option. These results suggest that the limb network does not exclusively pattern appendage outgrowth but is performing a more universal developmental function: radial patterning.

## INTRODUCTION

In the Metazoa, homologous networks of transcription factors are necessary for the development of some analogous structures in distantly related taxa. The limb patterning program is an example of this developmental process homology (Shubin et al., 1997; Erwin & Davidson, 2002; Pueyo & Couso, 2005). The limb program was first identified in the development of the proximal-distal axis of the *Drosophila* leg. The transcription factor *SP6-9/sp1* is upstream of other program members, *Dlx/dll, Pbx/exd, Meis/hth, Dac* and *Arx/ar*, each required for patterning specific regions of limb outgrowth (Panganiban et al., 1994; Panganiban et al., 1997; Dong et al., 2001, Dong et al., 2002; Peuyo & Couso, 2005; Estella et al., 2012; Campbell & Tomlinson, 1998). This network is necessary in both vertebrate and cephalopod limb development and is expressed in a similar proximodistal pattern in a diversity of outgrowths (Panganiban et al., 1997; Shubin et al., 1997; Maas & Bei, 1997; Mercader et al., 1999; Panganiban & Rubenstein, 2002; Prpic, 2003; Angelini & Kaufman, 2005; Pueyo & Couso, 2005; Shubin et al., 2009; Moczek & Rose, 2009; Capellini et al. 2011; Lapan & Reddien, 2011; Ibarretxe et al., 2012; Grimmel et al., 2016; Sanz-Navarro et al., 2019; Ramanathan et al. 2018; Setton & Sharma; 2018; Tarazona et al., 2019; Prpic, 2019). This suggests that, although each appendage is not homologous, an outgrowth program may have been present in the ancestor. Current fossil evidence and the prevalence of limbless taxa does not support an ancestor with appendages and therefore the network’s ancestral function remains unclear (Shubin et al., 1997; Erwin & Davidson, 2002; Pueyo & Couso, 2005). Many alternative hypotheses have been proposed, including an ancestral role in the nervous system, body axis formation and radial patterning (Minelli, 2000; Pueyo & Couso, 2005; Lemons et al. 2010; McDougall et al., 2011; Plavicki et al., 2016; Carroll et al., 1994; Erwin & Davidson, 2002). To understand the nature of this homology and how these co-option events occur, experiments with better sampling across the phylogeny of animals and greater diversity of developmental context are required.

Recent work identified a duplication of SP6-9 in cephalopods (McCulloch and Koenig, 2020). Both paralogs are expressed in the developing limb in the squid *Doryteuthis pealeii*, while one paralog, *DpSP6-9a*, shows unique expression in the lens-making cells during eye development (McCulloch and Koenig, 2020). With SP6-9 a known regulator in the limb patterning program, this new domain of expression could result in the co-option of the program in the cephalopod eye, providing a useful heterologous developmental context to better understand the network’s function.

The image-forming eye is a classic example of biological complexity and the lens is a requisite innovation in all high-resolution visual systems (Darwin, 1859; Arendt, 2009; Dakin, 1928; Walls, 1939; Koenig & Gross, 2020; Nilsson, 2013; Jonasova & Kozmik, 2008). Cephalopods have a single-chambered eye, morphologically convergent with the vertebrate eye, composed of a cup shaped retina and a single refractive lens (Packard, 1972). Here we perform the first in-depth molecular description of lens development in the squid *Doryteuthis pealeii*, we identify spaciotemporal expression of the limb patterning program in the developing eye and lens, and we demonstrate a negative regulatory role of canonical Wnt signaling upstream of the program.

## RESULTS AND DISCUSSION

### Cephalopod Lentigenic Cell Differentiation and Early Anterior Segment Heterogeneity

The anterior of the cephalopod eye, or the anterior segment, is composed primarily of lens generating cells (lentigenic cells) (Williams, 1909; Arnold, 1967; Brahma, 1978). Lentigenic cells are arranged circumferentially around the developing lens and extend long cellular processes, fusing into plates to form the lens (Figure 1A) (Meinertzhagen, 1990; Williams, 1909; Arnold, 1965; Arnold, 1967; West et al., 1995). We identified the first evidence of differentiated lentigenic cells starting at late stage 21, using a previously described nuclear morphology, unique to one of the three lentigenic cell types (LC2) (Figure 1B) (Arnold, 1967; West et al., 1995; Koenig et al., 2016). The number of LC2 cells continues to grow until reaching pre-hatching stage (Stage 29). We performed staged *in situ* hybridization for a homolog of *DpS-Crystallin*, the most abundant family of proteins in the cephalopod lens (Chiou, 1984; West et al., 1994) (Supplemental Figure 1). The first evidence of expression corresponds to changes in nuclear morphology at stage 21 (Figure 1C).

**Figure 1:**
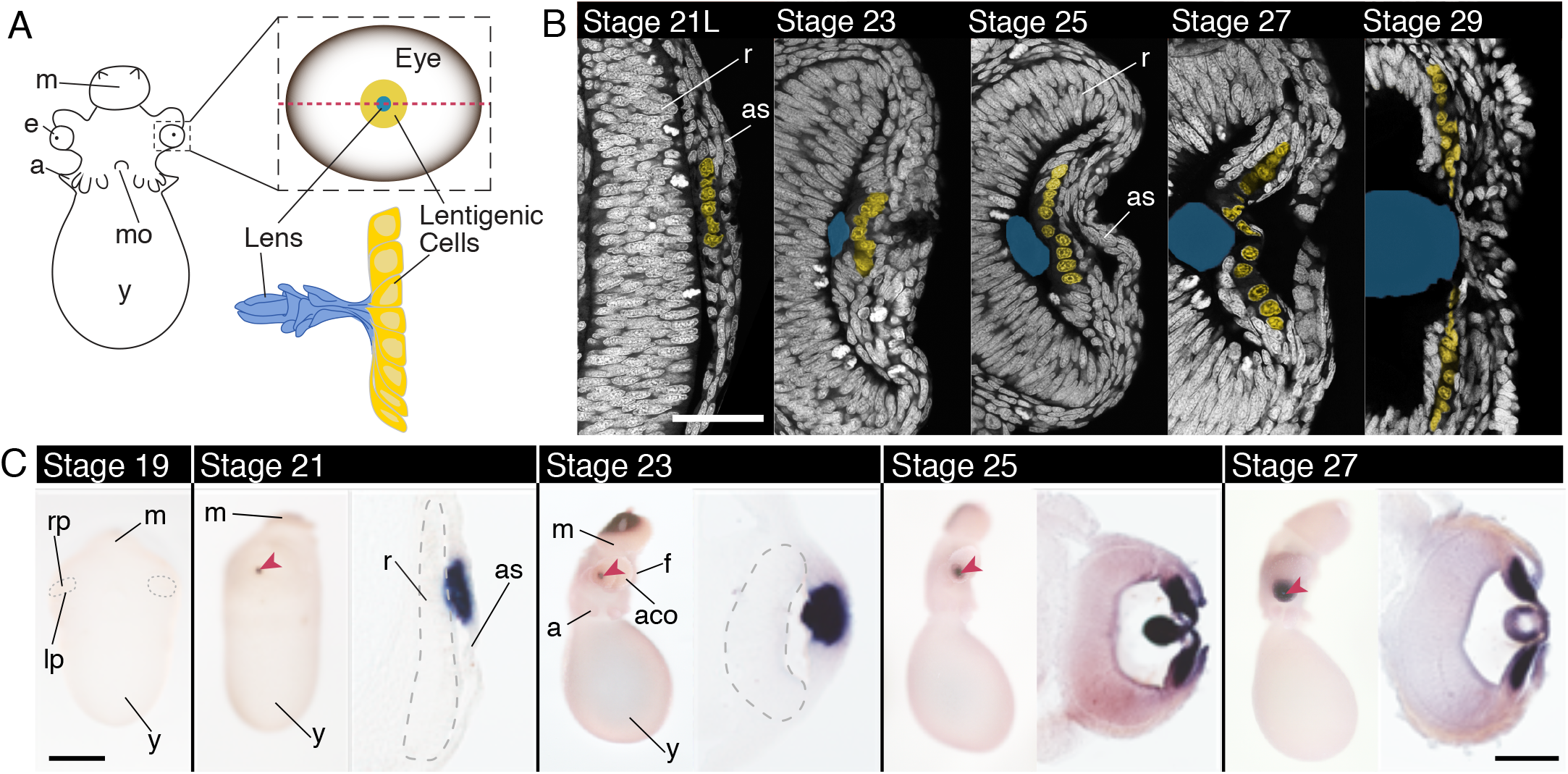
Lentigenic cell differentiation **and *DpS-Crystallin* expression in the squid**. A) Cartoon diagram of a squid embryo (anterior), en face cartoon of the developing eye (red dotted line shows cross-section plane) and developing lentigenic cells and lens. (Cartoon of lens and lentigenic cells based on Arnold, 1967) B) Cross-section of the developing anterior segment at Arnold stages 21 late, 23, 25, 27 and 29 identifying differentiation of lentigenic cells (Arnold, 1968). White: Sytox-Green labeling nuclei, Yellow: False-colored lentigenic cell nuclei corresponding to the LC2 population identified by nuclear morphology (Arnold, 1967; West et al., 1995; Koenig et al., 2016). Blue is the outline of the lens, as identified using phalloidin staining (not shown). First evidence of LC2 cells is late stage 21. Lentigenic cell number multiplies and distribution grows across the anterior segment (*as)* throughout development. Scale is 50 microns. C) *In situ* hybridization of *DpS-Crystallin* in whole-mount and cryo-section. Stage 19 is an anterior view, the boundary between the retina placode and the lip cells is highlighted with a dotted line. No *DpS-Crystallin* expression is apparent at this stage. Stage 21-27 are shown in a lateral view of the embryo on the left and a cross-section of the eye on the right. Anterior of the embyro is down in the sections. The retina is outlined with a dashed grey line in stage 21 and 23. *DpS-Crystallin* expression corresponds with LC2 lentigenic cell population. Scale is 500 microns in whole mount images. Scale is 100 microns in sectioned images. *as*, anterior segment; *a*, arm; *aco*, anterior chamber organ; *e*, eye; *f*, funnel *lp*, lip; *m*, mantle; *mo*, mouth; *rp*, retina placode; *r*, retina; *y*, yolk. Red arrow highlights the lens.

We sought to understand the molecular heterogeneity of cells in the early developing anterior segment, of which nothing is currently known. Using previously published candidates and RNA-seq data, we performed *in situ* hybridization screens at stage 23 to identify unique cell populations (Koenig et al., 2016; Ogura et al., 2013). We find *DpSix3/6* at stage 23 expressed in the anterior segment in the distal cells that make a central cup (*cc*), as well as a marginal population of cells in the most proximal tissue (*pm*) (Figure 2B’’). The proximal central cells lacking *DpSix3/6* expression correspond to the LC2 population (Figure 2A’’ &B’’). Asymmetry along the animal anterior-posterior axis in the eye is also apparent, with enrichment on the anterior side of the animal (Figure 2B’’). We also find the gene *DpLhx1/5*, expressed in a distal-marginal population of cells in the anterior segment (*dm*), and excluded from the distal central cup cells (*cc*) (Figure 2C’’). Together these genes show distinct populations of cells present early in development and provide a helpful molecular map of the anterior segment tissue at this time point: central cup cells (*cc*), LC2 cells (*lc2)*, proximal-marginal cells (*pm)*, and distal-marginal cells (*dm*) (Figure 2).

**Figure 2:**
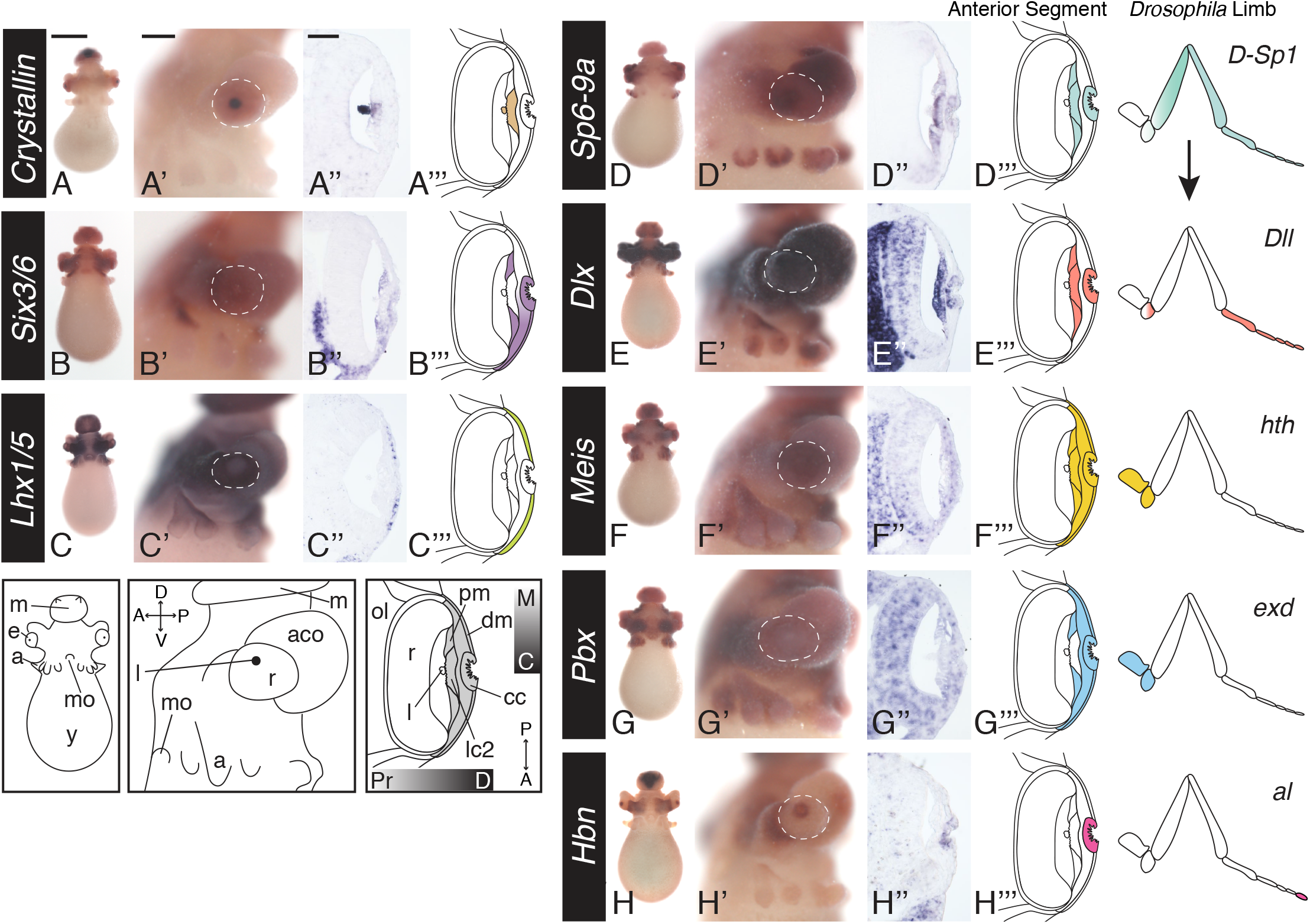
Limb patterning program expressed in the developing anterior segment. For each gene: left to right, anterior whole-mount view, lateral whole-mount view (anterior left), cross-section (anterior is down), cartoon summary of anterior segment expression. Dotted white outline in lateral view outlines the perimeter of the eye. A-C) Defining cell populations in the developing anterior segment at stage 23. A, A’, A’’) *DpS-Crystallin* expression in the anterior segment at stage 23, expressed in the proximal, central cells corresponding with the LC2 cells (*lc2*). Expression is also apparent in the lens. B, B’, B’’) Expression of *DpSix3/6*. B’’) Expression is apparent in the distal, central cup cells (*cc*) and the proximal-marginal (*pm*) anterior segment cells. The proximal-central cells (*lc2*) lack expression of *DpSix3/6*. C, C’, C’’) *DpLhx1/5* expression. C’’) Expression of *DpLhx1/5* is found in the distal-marginal cell (*dm*) population. Expression is excluded from the central cup (*cc*). D-G) Expression of the limb patterning program genes. Summary of the proximodistal expression of each *Drosophila* homolog during proximodistal patterning of the limb is shown on the right H) Prd-like homolog *Homeobrain* (*Hbn*) expression in the distal, central cup cells. *a*, arms; *aco*, anterior chamber organ; *cc*, cup cells; *dm*, distal-marginal cells; *e*, eye; *l*, lens; *lc2*, LC2 cells; *m*, mantle; *mo*, mouth; *pm*, proximal-marginal cells; r, retina; *y*, yolk. Anterior segment highlighted in grey in the cartoon. Orientation abbreviations: M, marginal; C, central; Pr, proximal; D, Distal; A, anterior; P, posterior. Scale for whole-mount anterior view is 500 microns. Scale for lateral whole-mount view 200 microns. Scale for sectioned images 50 microns.

### Proximal-Distal Limb Patterning Genes in the Anterior Segment of the Cephalopod

To assess whether genes involved in appendage patterning may be required for cephalopod lens development, we identified and performed *in situ* hybridization for the genes *Dlx, Pbx, Meis*, and *Dac* at stage 21 and 23 (Figure 2, Supplemental Figure 2). All genes were clearly expressed in the developing anterior segment and lentigenic cells with the exception of *DpDac* (Figure 2E-G, Supplemental Figure 2I-2J’). We find *DpDlx* and *DpSP6-9a* have overlapping expression, in the central cup cells (*cc*) and all proximal cells (LC2 and *pm*) (Figure 2D-E’’’). *DpPbx* and *DpMeis* are both broadly expressed in the anterior segment during lens development, with *DpPbx* excluded from the LC2 cells (Figure 2F’’& 2G’’).

It is known that the transcription factor *aristaless* is necessary for the most distal tip of the *Drosophila* limb in the limb program (Campbell and Tomlinson, 1998). The evolutionary relationship of Prd-like homologs (Arx/Aristaless, Alx/Aristaless-like, Rx/Retinal Homeobox and Hbn/Homeobrain) is ambiguous across species (Schiemann et al., 2017). We identified three candidate Prd-like genes in *D. pealeii* and performed *in situ* hybridization for all three homologs, *DpHbn, DpPrdl-1* and *DpPrdl-2* (Supplemental Figure 2K, L) (Koenig et al, 2016). *DpHbn* is expressed in the anterior segment in the distal central cup cells (*cc*) while *DpPrdl-1* and *DpPrdl-2* are excluded from the eye (Figure 2H’’ and Supplemental Figure 2C, C’, K and L). *DpHbn’s* central, distal expression recapitulates *aristaless* expression in the developing *Drosophila* limb.

Our data show that the majority of the proximal-distal patterning genes in the developing limb, including *SP6-9, Dlx, Meis, Pbx*, as well as the Prd-like homolog, *Hbn*, show expression in concentric and overlapping cell populations surrounding the developing lens in the squid (Figure 2). This pattern of expression is strikingly similar to the bullseye-like pattern of expression of these genes in the developing *Drosophila* limb imaginal disc and suggests a co-option of this regulatory program for a new function: patterning the cephalopod anterior segment and lens (Angelini & Kaufman, 2005).

### Canonical Wnt Signaling Genes Expressed During Anterior Segment Development

The duplication of SP6-9 in cephalopods provides a substrate for the evolution of cis-regulation, which could result in novel expression of the limb patterning program in the cephalopod lens. In appendage outgrowth, active Wnt signaling is upstream of the expression of SP6-9 (Cohen, 1990; Estella et al., 2003). To assess whether Wnt may be acting upstream in the cephalopod anterior segment or whether novel regulatory mechanisms may be at play, we performed *in situ* hybridization for members of the Wnt signaling pathway at stage 21 and stage 23 (Figure 3, Supplemental Figure 3). We were interested in identifying cells in the anterior segment or in adjacent tissue that may be a source of the Wnt morphogen. We performed *in situ* hybridization for seven Wnt homologs, with most *Wnt* genes expressed in the retina (Figure 3A’, 3C’, 3D-G). *DpWnt8, DpWnt11* and *DpProtostome-specific Wnt* show the most robust retinal expression (3A’, 3F & 3G) and *DpWnt7* is the only Wnt expressed in the anterior segment (Figure 3C). *DpWnt6* showed no evidence of expression in the developing eye (data not shown). These data support the hypothesis that Wnt signals emanating from neighboring tissues could regulate anterior segment development.

**Figure 3:**
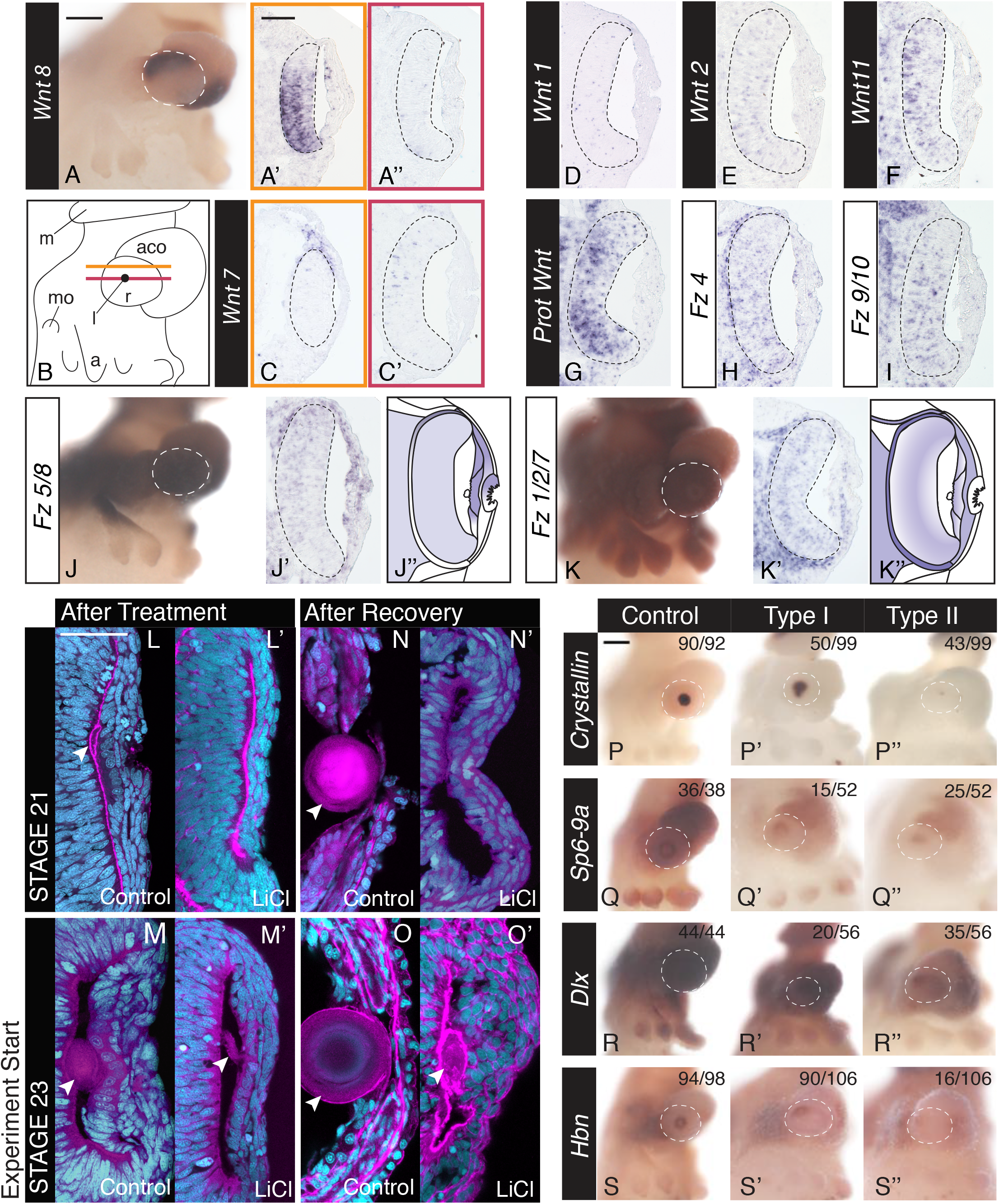
Wnt signaling pathway expression in the developing cephalopod eye. A-G) *Wnt* gene expression at stage 23. Based on expression, Wnt7, Wn8, Wnt2, Wnt11 and Prot Wnt are possible candidates to signal the anterior segment. A) Lateral, whole-mount expression of *Wnt8*. A’) Dorsal retina expression of *Wnt8*. Location of the section indicated by the orange line in B. A’’) Central section lacking retina expression. Location of the section indicated by the red line in B. B) Cartoon of the lateral whole-mount embryo at stage 23. Orange and red lines correspond to the location of the two sections shown in A, A’, and C, C’. D-G) Expression of other *Wnt* homologs in central sections. H-K) Expression of Frizzled receptors at stage 23. Fz1/2/7 shows asymmetric expression and Fz5/8 shows specific exclusion from the central cup cells. J and K are lateral view of the whole mount expression. J’’ and K’’ are cartoons of expression in J’ and K’ respectively. Black dotted line in sectioned images show the perimeter of the retina. L-O) Anterior segment and lens morphology after Wnt agonist treatment (LiCl). Embryos were cryosectioned and stained with sytox-green (nuclei, cyan) and phalloidin (F-actin, magenta).. L and L’) Control and LiCl agonist treatments started at stage 21, treated for 24 hours and fixed immediately. M and M’) Control and Wnt agonist (LiCl) treatments started at stage 23 for 24 hours and fixed immediately. N and N’) Control and Wnt agonist (LiCl) treatments started at stage 21, treated for 24 hours and allowed to recover for 48 hours and fixed. O and O’) Control and Wnt agonist (LiCl) treatments started at stage 23, treated for 24 hours and allowed to recover for 48 hours and fixed. Arrowhead highlights the lens. P-S) *In situ* hybridization of anterior segment markers after 24 hour control and LiCl treatments starting at stage 23. Phenotypes are characterized as Type I (mild) and Type II (severe). The white dotted line outlines the eye in the lateral image and the number of eyes scored in control and the two phenotypes is found in LiCl treated animals in the top right corner. Scale for all lateral whole-mount view images is 200 microns. Scale for all sectioned images is 50 microns. Anterior is down in all sectioned images. White dotted line in whole mount images identify the perimeter of the eye. *m*, mantle; *a*, arms; *aco*, anterior chamber organ; *mo*, mouth; *r*, retina; *l*, lens.

To identify cells with potential active Wnt signaling, we analyzed the expression of Fz genes, which encode a family of Wnt receptors. We find that *DpFz* receptors are expressed broadly throughout the embryo. A subset of these (e.g. *DpFz1/2/7, DpFz4*, and *DpFz5/8*) are expressed in the majority of cells in the anterior segment, while others, like *DpFz9/10*, are excluded from the anterior segment (Figure 3H-K, Supplemental Figure 3). On close examination we find that *DpFz5/8* is excluded asymmetrically in the anterior segment and may be important for anterior-posterior patterning (Figure 3J’ & J’’). *DpFz1/2/7* is excluded from the distal-marginal cells and central cup cells and interestingly, the central cup cells lacking *DpFz1/2/7* are those that express all the limb patterning program genes (Figure 3K’&K’’). These data suggested that the exclusion of active Wnt signaling may be important in the cephalopod anterior segment, supporting a potential negative regulatory role for Wnt signaling.

### Ectopic Wnt Activation Leads to the Loss of the Lens

To assess the hypothesis that Wnt signaling is playing a negative regulatory role in anterior segment development, we utilized well-characterized pharmacological compounds that act as agonists of the Wnt pathway (Hedgepeth et al. 1997; Klein & Melton; Sato et al., 2004). We empirically determined a working concentration of both LiCl (0.15M) and CHIR99021 (250um). We bathed embryos in the compound or vehicle control for 24 hours at stage 21, the onset of lentigenic cell differentiation, and immediately fixed thereafter. Embryos were sectioned and assessed for phenotypes. Stage 21 control embryos show a thickened anterior segment, identifiable lentigenic cells, and small lens primordia (Figure 3L). LiCl-treated stage 21 embryos show a complete absence of lens formation: No anterior segment thickening, lentigenic cells, or lens tissue. These data suggest that ectopic Wnt pathway activation inhibits lens and anterior segment development (Figure 3L’, Supplemental Figure 4A). CHIR99021 treatment showed similar phenotypes (Supplemental Figure 4A). We assessed LiCl treated and control animals for cell death and find little difference between control and treated eyes suggesting that toxicity is unlikely the reason for these phenotypic changes (Supplemental Figure 4B).

**Figure 4:**
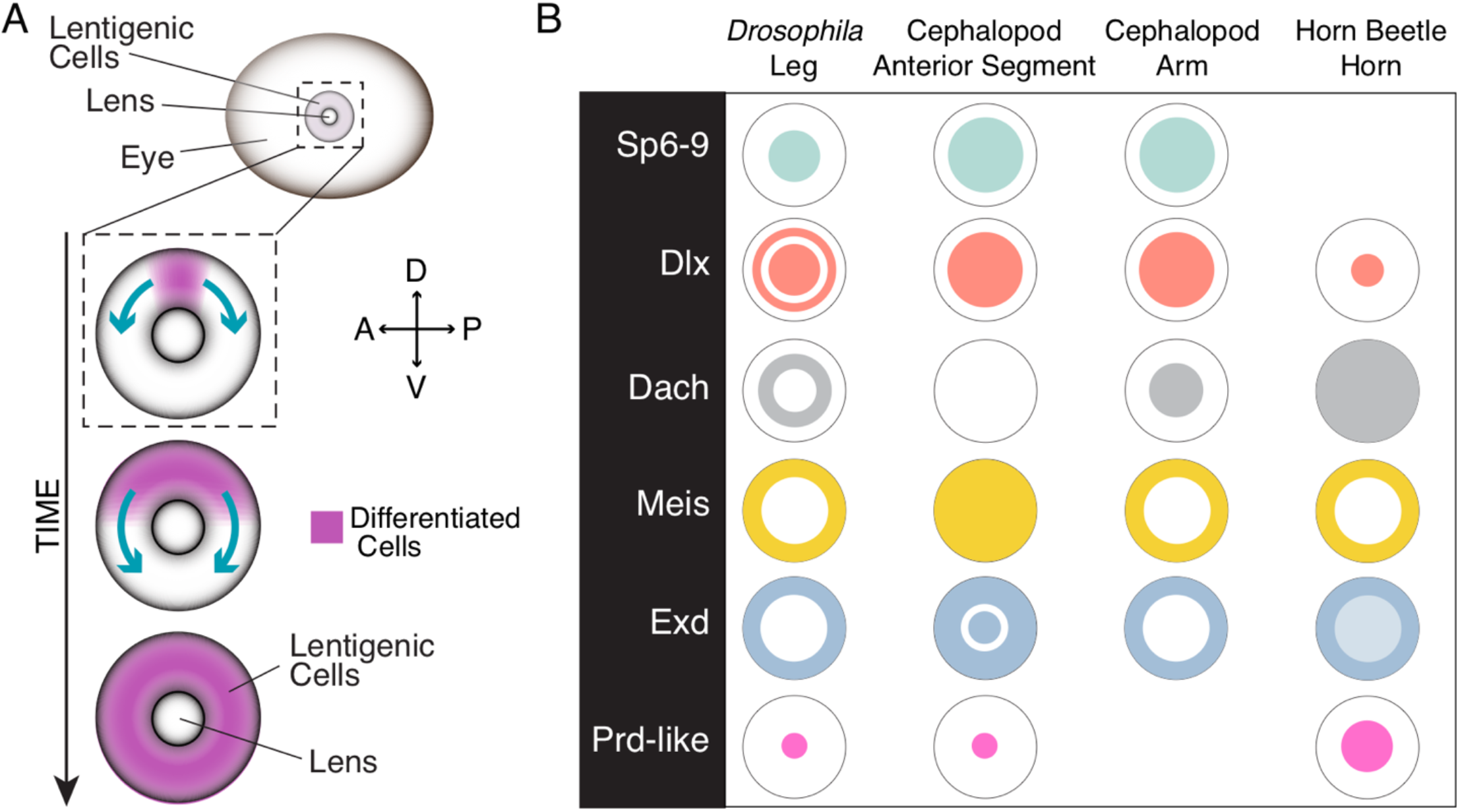
Ectopic Wnt signaling activation leads to loss of the lens. A) Model for lentigenic cell differentiation at stage 21. LC2 lentigenic cells differentiate on the dorsal side of the eye first, with a wave moving ventrally. Type I *DpS-Crystallin* embryos have been interrupted in progress. B) En face summary of sample radial expression of the limb patterning program across developmental contexts (Tarazona, 2019; reviewed in Moczek, 2009 and Angelini & Kaufman, 2005).

We were interested in the consequence of activating the Wnt pathway after lentigenic cell differentiation. We performed the same 24 hour LiCl exposure at stage 23 and find the lens smaller and the anterior segment less thick than control animals, but lentigenic cells and lens tissue remain identifiable. This suggests that ectopic Wnt signaling does not impact cell identity in differentiated lentigenic cells (Figure 3M & M’).

The lack of lens growth in stage 21 treated animals may be a result of an imposed delay in lens formation or it may be a result of the loss of lens potential. To differentiate between these possibilities we allowed treated animals to recover. We bathed experimental and control embryos, at both stage 21 and 23, for 24 hours, washed out the solution and allowed animals to develop for an additional 48 hours. LiCl treated stage 21 embryos never recover a lens (Figure 4N & 4N’) while LiCl treated stage 23 embryos do form a small but morphologically abnormal lens (Figure 4O & 4O’). This abnormal lens is larger than the lens found in animals immediately fixed after treatment, suggesting that existing lentigenic cells at stage 23 continue to contribute to lens formation and growth. However, because the stage 23 treated lens is markedly smaller than control, it suggests that further lentigenic cell differentiation is lost in treated animals. These data suggest that ectopic Wnt signaling leads to the disruption of lens potential and the lack of proper lentigenic cell differentiation.

Despite the remarkable loss of the lens, these data do not clearly distinguish between the loss of lentigenic cell fate or proper cell function, such as the growth of the cellular processes that form the lens. To assess if lentigenic cell fate is lost, we performed *in situ* hybridization experiments for *DpS-Crystallin* on LiCl treated animals. We saw two types of expression phenotypes, either a significant decrease (Type I) or a complete loss (Type II) in *DpS-Crystallin* expression as compared to control (Figure 4P, P’ & P’’). We find all *DpS-Crystallin* expression exclusively dorsal to the site of lens formation suggesting that these cells may differentiate first. These data show that ectopic Wnt signaling results in the loss of lentigenic cell fate and that our treatment may have interrupted a dorsal-to-ventral wave of differentiation in some embryos (Figure 4A). In addition, we assessed other anterior segment markers, including *DpSix3/6* and *DpLhx1/5*, and these genes show a consistent loss of expression in the most severe phenotypes, (Supplemental Figure 4C).

### Limb Patterning Program Regulatory Evolution

To address if Wnt signaling is upstream of the limb patterning program, we performed *in situ* hybridization of limb transcription factors after LiCl treatment (Figure 3Q-3S, Supplementary Figure 4C). Similar to *DpS-Crystallin* expression, we again see a mild reduction (Type I) or loss and severe reduction (Type II) of expression. Our milder phenotypes, again, show a dorsal asymmetry, which can be most easily seen in *DpSP6-9A, DpDlx* and *DpHbn* (Figure 3Q, Q’, Q’’, 3R, R’, R’’ and 3S, S’, S’’). Changes are also visible but less obvious in *DpPbx* and *DpMeis* expression, with *DpPbx* only showing a mild phenotype (Supplemental figure 4C). These data support the placement of Wnt signaling upstream of the limb patterning program in a negative regulatory role.

### Conclusion

Our findings indicate that the limb patterning program has been co-opted for anterior segment and lens development in cephalopods and that this co-option does not have a homologous upstream regulatory relationship with Wnt signaling as found in the limb (Estella et al., 2003; Tarazona et al., 2019). This change in signaling and the known duplication of SP6-9 identifies the paralog SP6-9a as a mediator of limb patterning program co-option in the anterior segment. Finally, with little similarity between limb and lens, our work suggests that the function of the limb patterning program in a limbless ancestor was likely a more generic developmental function than outgrowth. Considering present findings, previous work and hypotheses we conclude that the ability to pattern in a radial fashion, as previously proposed, is a more inclusive and likely ancestral function (Figure 4B) (Carroll et al., 1994; Erwin & Davidson, 2002). This work shows the cephalopod lens to be a unique context for future investigation of comparative regulatory changes responsible for co-option, and for identifying the regulatory mechanisms responsible for the emergent radial pattern found in embryos across species.

## Methods

### Animal Husbandry

*Doryteuthis pealeii* egg sacks were obtained from the Marine Biological Labs. Egg sacks were kept at 20 degrees Celsius. Although not required, European guidelines for cephalopod research were followed.

### Histology and TUNEL Staining

Embryos were fixed at 4 degrees Celsius overnight in 4% PFA in filter-sterilized seawater. After fixation embryos were transitioned into 15% and 30% sucrose and embedded in TFM and stored at -80 degrees Celsius. Embryos were cryosectioned in 12 um sections, stained with Sytox Green 1:1000 and Phalloidin 555 1:300 in PBS overnight (Molecular Probes). Tunel stained tissue was processed after sectioning using the Click-iT TUNEL Alexa Fluor 488 kit according to manufacturer’s instructions (Invitrogen). Embryos were mounted in VECTASHIELD Hardset antifade mounting medium and imaged on a Zeiss 880 confocal.

### Homolog Identification and Phylogenetics

Genes were preliminarily identified using reciprocal BLAST with *Mus musculus* and *Drosophila melanogaster* sequences as bait with the exception of S-Crystallin where previous *Doryteuthis opalescens* sequences were also used (Altschul et al., 1990). Top hits in the *D. pealeii* transcriptome were trimmed for coding sequence and translated to amino acid sequences. To find related sequences, BLASTp was used, searching only the RefSeq protein database in NCBI filtered for vertebrate and arthropod models, as well as spiralian models for when published annotated sequences could be found. The top hits of each gene name were downloaded and aligned with our *D. pealeii* sequences for each tree using MAFFT in Geneious (Katoh, 2002). To check sequence redundancy and proper outgroups quick trees were made using FastTree. We constructed maximum-likelihood trees on the FASRC Cannon cluster supported by the FAS Division of Science Research Computing Group at Harvard University (Price et al. 2010). Using PTHREADS RAxML v.8.2.10, we ran the option for rapid bootstrapping with search for best maximum likelihood tree, resampling with 1000 bootstrap replicates, the PROTGAMMAAUTO model of amino acid substitution, and otherwise default parameters (Stamatakis, 2014). Fasta alignments, Nexus tree files are found in the Supplemental Data Folder. All PDF versions of the trees are found in Supplemental Figure 1.

### Cloning and Probe Synthesis

Embryos stg 21-29 were crushed in Trizol reagent. RNA was extracted using standard phenol-chloroform extraction with a clean-up using the Qiagen RNeasy Micro kit. cDNA was synthesized using iScript (Bio-Rad) according to manufacturer protocols. Primers were designed using Primer3 in the Geneious software package from available transcriptomic data (Koenig et al., 2016). PCR products were ligated into the Pgem-T Easy plasmid and isolated using the Qiagen miniprep kit. Plasmids were linearized using restriction enzymes. Sense and anti-sense probes were synthesized using T7 and SP6 polymerase with digoxygenin labelled nucleotides.

### *In situ* Hybridization

Embryos were fixed as previously described (Koenig et al. 2016) and were dehydrated in 100% ethanol and stored at -20 degrees Celsius. Whole-mount *in situ* hybridization was performed as previously described (Koenig et al., 2016). Embryos were imaged using a Zeiss Axio Zoom.V16. Embryos were fixed for sectioning overnight in 4% PFA in artificial seawater and dehydrated in 100% ethanol. Embryos were transitioned into histoclear and embedded in paraffin. Embryos were sectioned on a Leica RM2235 microtome in 5-micron sections. Sections were dewaxed for *in situ* in Histoclear, rehydrated through an EtOH series, and re-fixed for 5 minutes at 4 degrees Celsius in 4% PFA in PBS. Embryos were exposed to Proteinase K for 20 minutes at 37 degrees Celsius and then quenched with glycine. The embryos were then de-acetylated with acetic anhydride. Slides were then pre-hybridized at 65 degrees Celsius for 30-60 minutes and then exposed to probe overnight. Slides were washed in 50% formamide/1x SSC/0.1% Tween-20 hybridization buffer twice, then twice in 1x SSC, .2x SSC and 0.02x SSC, all at 70 degrees Celsius. The slides were then washed at room temperature in MABT three times and blocked in Roche Blocking Buffer for an hour. Slides were incubated in Anti-Dig antibody (Roche) at 1/4000 overnight at 4 degrees Celsius. Slides were washed with MABT and then placed in AP reaction buffer. Slides were then exposed to BCIP/NBT solution until reacted and stopped in PBS. Slides were counterstained with Sytox 1:1000 overnight. Slides mounted in ImmunoHistoMount (Abcam) and imaged on a Zeiss Axioscope. *DpS-Crystallin* embryo *in situs* were transitioned to sucrose and embedded after imaging in whole-mount. Embryos were image on a Zeiss Axioscope.

### *Ex ovo* Experimental Culture

*Ex ovo* culture was performed as previously described in Koenig, 2016. Embryos were bathed in .25 M, .15 M and .07 M LiCl and 100nm, 250nm and 500nm concentration of Wnt Agonist (CHIR99021) in Pen-Step filter-sterilized seawater to determine a working concentration. Control animals were bathed in equivalent amounts of DMSO or Pen-Strep alone.

## Supplemental Data Files

### RAxML Maximum Likelihood trees, 1000 bootstraps

ANTP_ML_1000bs_final.nex

Axin_ML_1000bs_final.nex

Cry_ML_1000bs_final.nex

Dach_ML_1000bs_final.nex

Dsh_ML_1000bs_final.nex

Fz_ML_1000bs_final.nex

GSK3_ML_1000bs_final.nex

Lhx_ML_1000bs_final.nex

LRP1_ML_1000bs_final.nex

Pangolin_ML_1000bs_final.nex

Prd_domain_ML_1000bs_final.nex

TALE_ML_1000bs_final.nex

Wnt_ML_1000bs_final.nex

### MAFFT sequence alignments

ANTP_ML_1000bs_final.fasta

Axin_ML_1000bs_final.fasta

Cry_ML_1000bs_final.fasta

Dach_ML_1000bs_final.fasta

Dsh_ML_1000bs_final.fasta

Fz_ML_1000bs_final.fasta

GSK3_ML_1000bs_final.fasta

Lhx_ML_1000bs_final.fasta

LRP1_ML_1000bs_final.fasta

Pangolin_ML_1000bs_final.fasta

Prd_domain_ML_1000bs_final.fasta

TALE_ML_1000bs_final.fasta

Wnt_ML_1000bs_final.fasta

## Authors’ contributions

K.M.K. designed the experiments. S.N., K.J.M., F.N., C.D., J.C., and K.M.K. performed experiments. K.J.M. performed phylogenetic analyses. K.M.K., S.N., and K.J.M. wrote the manuscript with consultation from all authors.

## Competing Interests

Authors declare no competing interests.

## Funding

This work is supported by the Office of the NIH Director 1DP5OD023111-01, and the John Harvard Distinguished Science Fellowship to K.M.K.

## Acknowledgements

The authors would like to thank the Koenig and Srivastava lab members for helpful discussions as well as Kevin Woods and the John Harvard Distinguished Science Fellows community for support. We would like to thank Jeffrey Gross, Alex Schier, Mansi Srivastava, and Andrew Murray for comments on the manuscript. We also thank the Marine Biological Labs, the Marine Resources Center, Owen Nichols, Ernie Eldredge, and Shannon Eldredge for assisting in the acquisition of embryos. We would also like to acknowledge the Harvard College undergraduates of LS50: Integrated Science Laboratory Course: Zach Alerte, Vlad Batagui, Eli Burnes, Stephen Casper, Chris Chen, Ahab Chopra, Ralph Estanboulieh, Lily Gao, Pedro Garcia, Saimun Habib, Harry Hager, Maxwell Ho, Charlie Horowitz, Ray Jiang, Prashanth Kumar, Truelian Lee, Arian Mansur, Matthew Mardo, Mark Theodore Meneses, Kendrick Nguyen, Francesco Rolando, Simon Schnabl, Taylor Shirtliff-Hinds, Sorscher Lincoln, William Stainier, Avi Swartz, David Szanto, Sophia Tang, Joey Toker, Analli Torres, Nina Uzoigwe, Rowen VonPlagenhoef, Evelyn Wong, Alexandra Zaloga, Maxwell Zhu.

## Supplemental Tables

**Supplemental Table 1:**
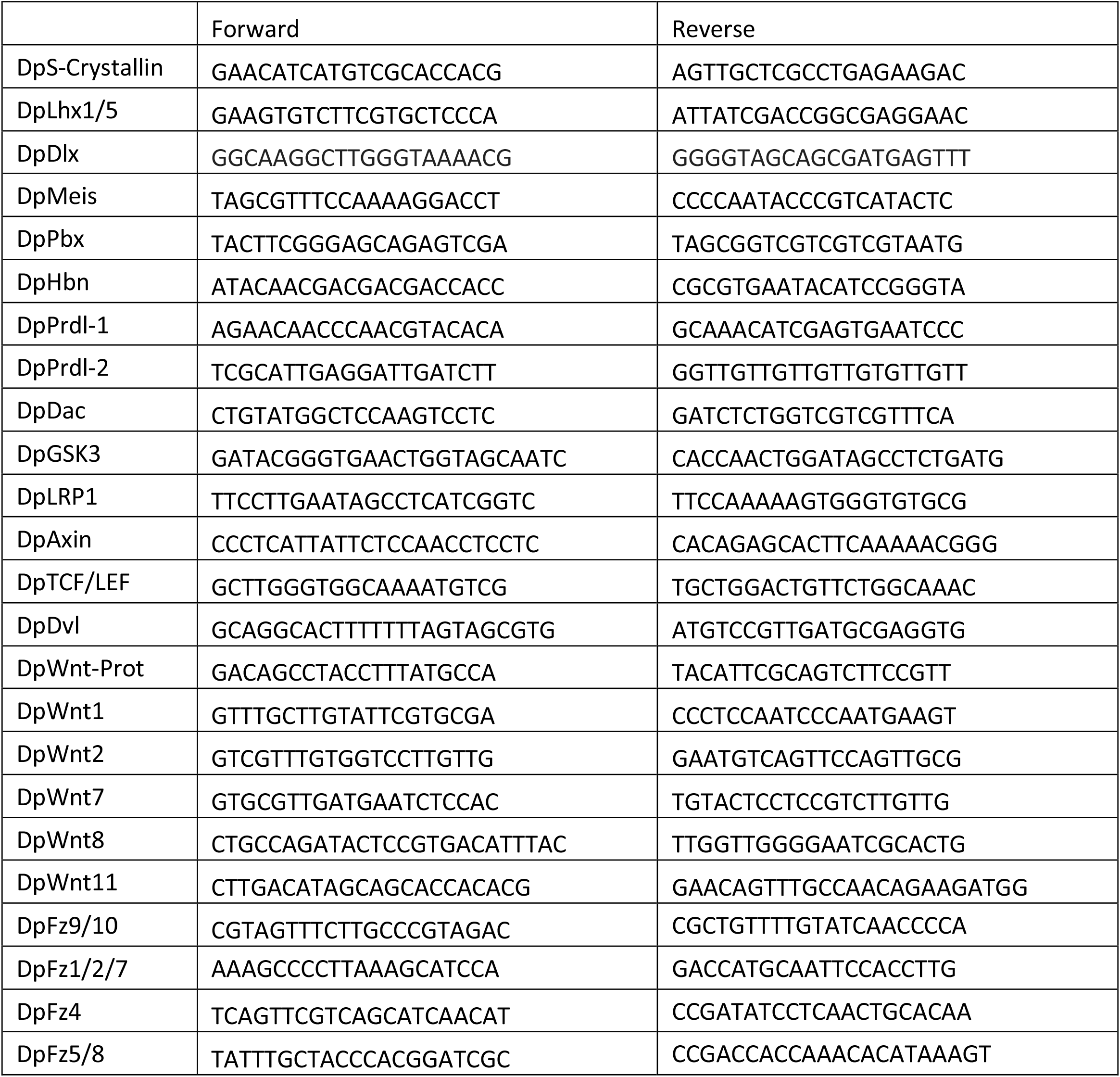
All Primer sequences

## Supplemental Figure titles

**Sup Figure 1:**
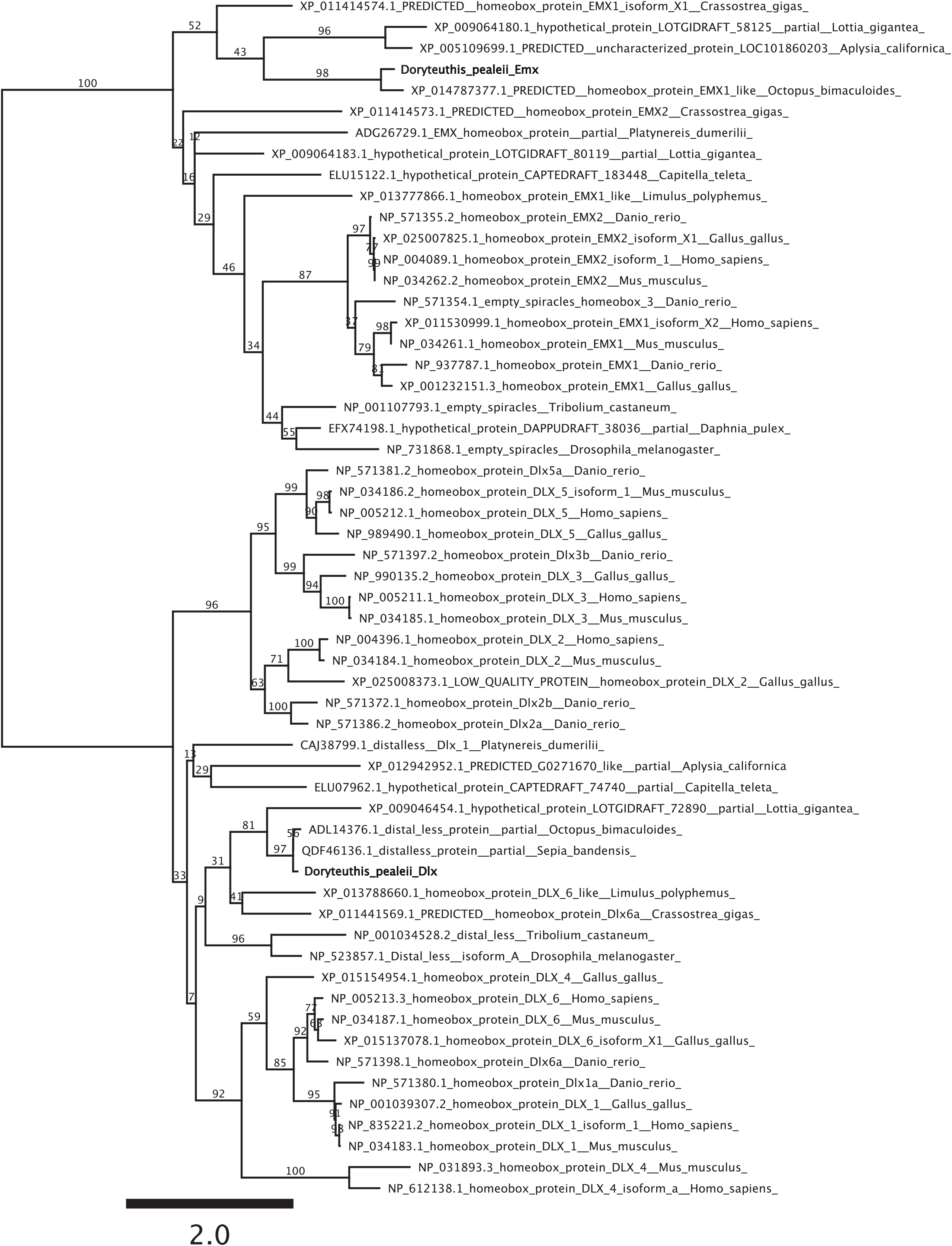
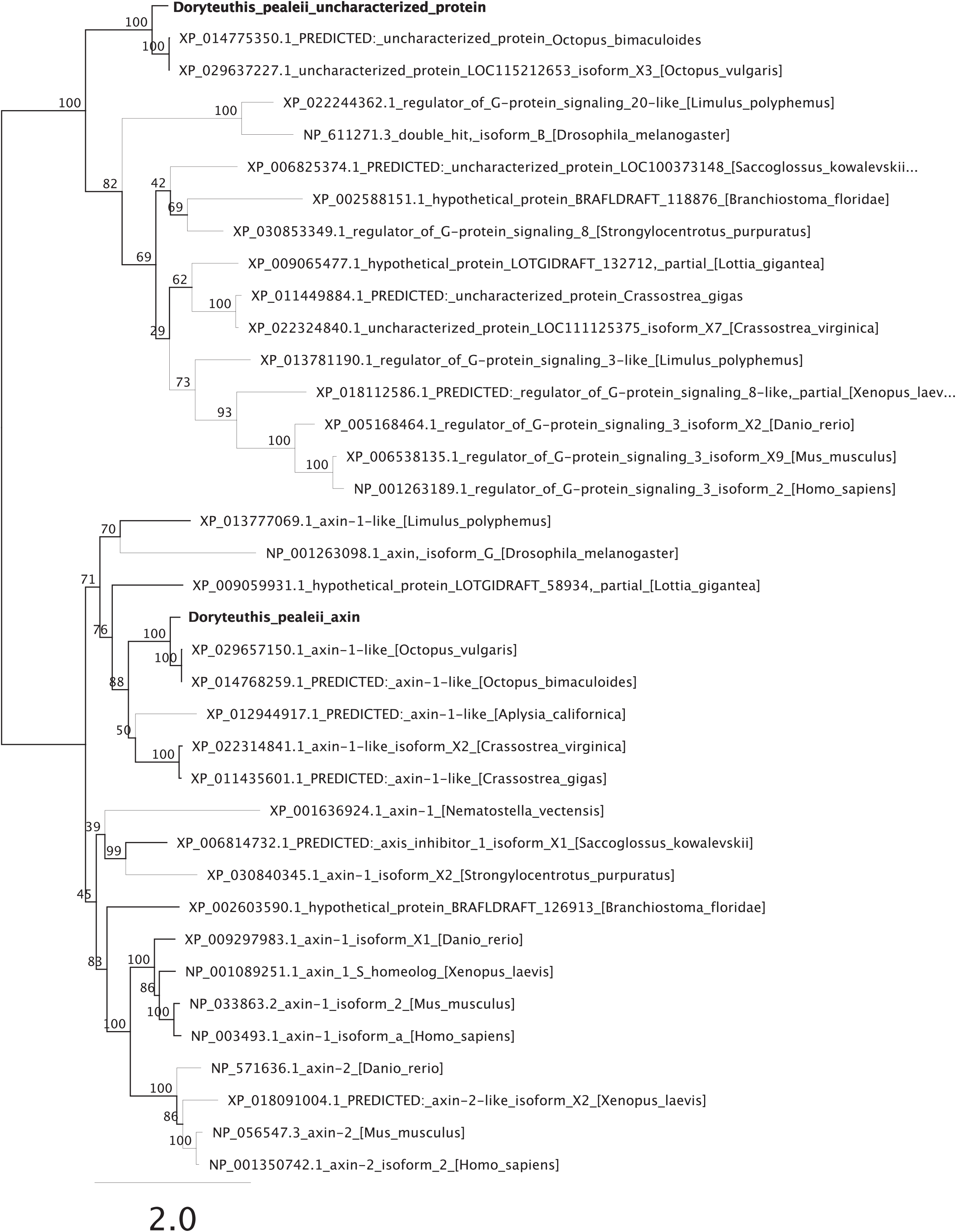
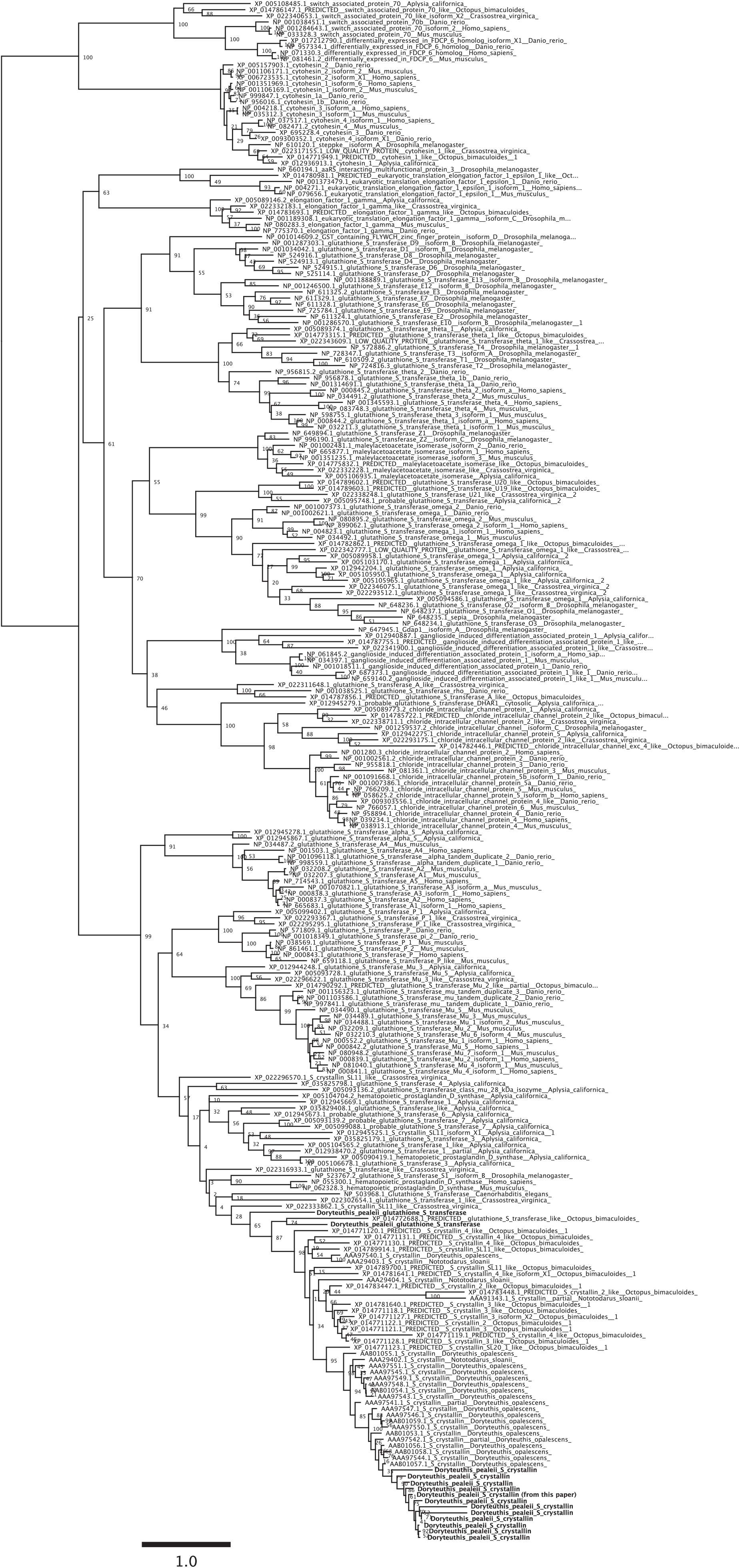
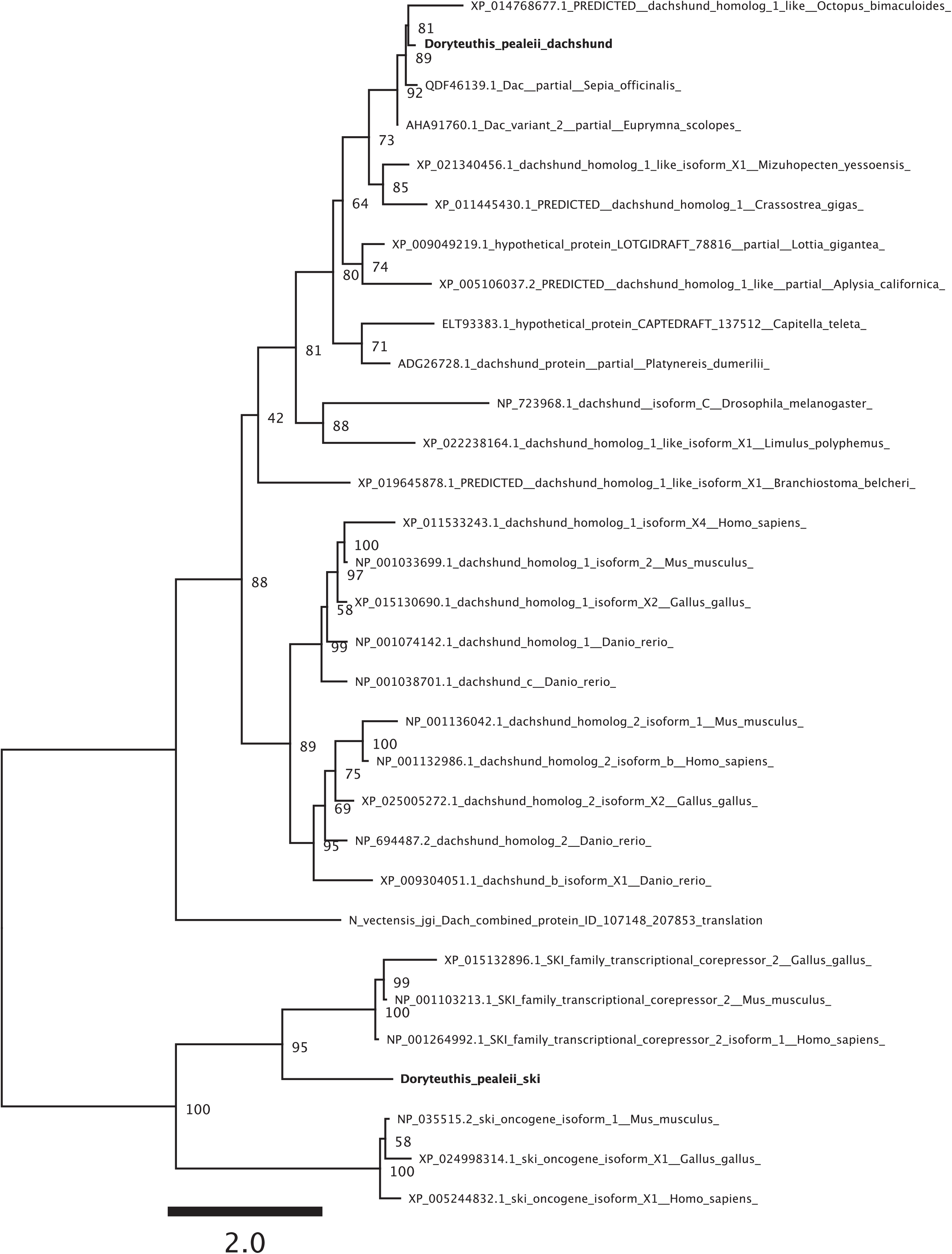
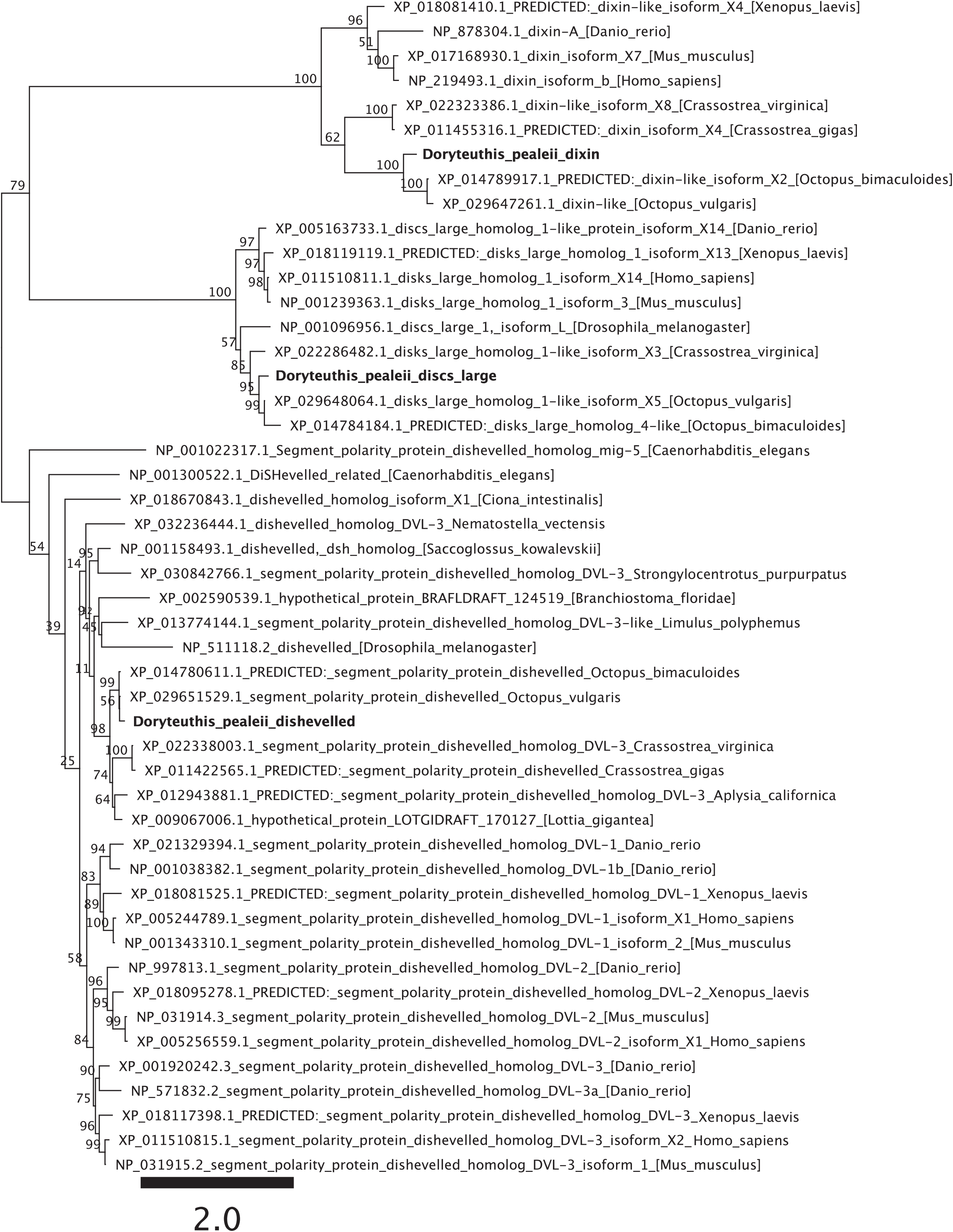
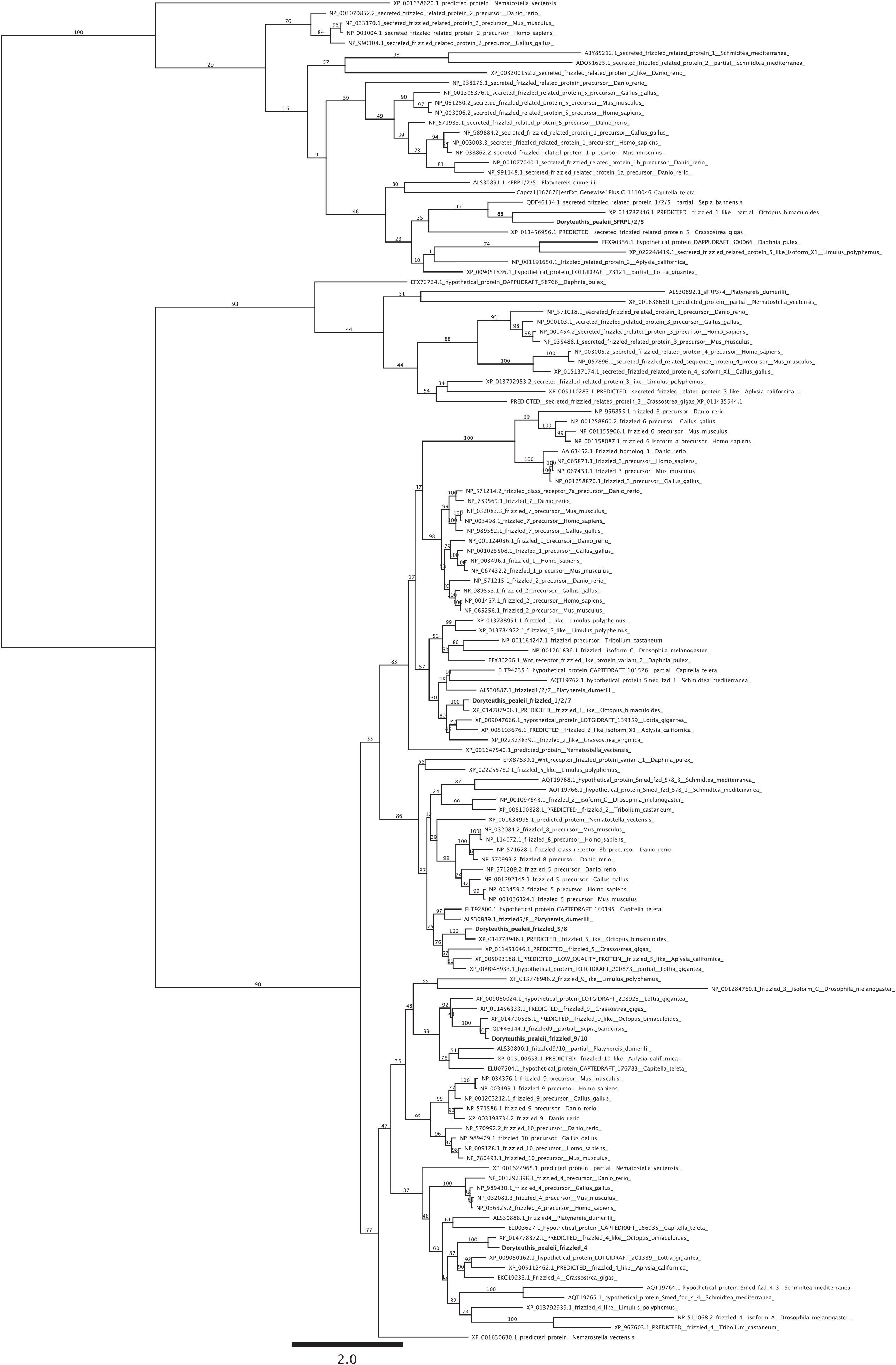
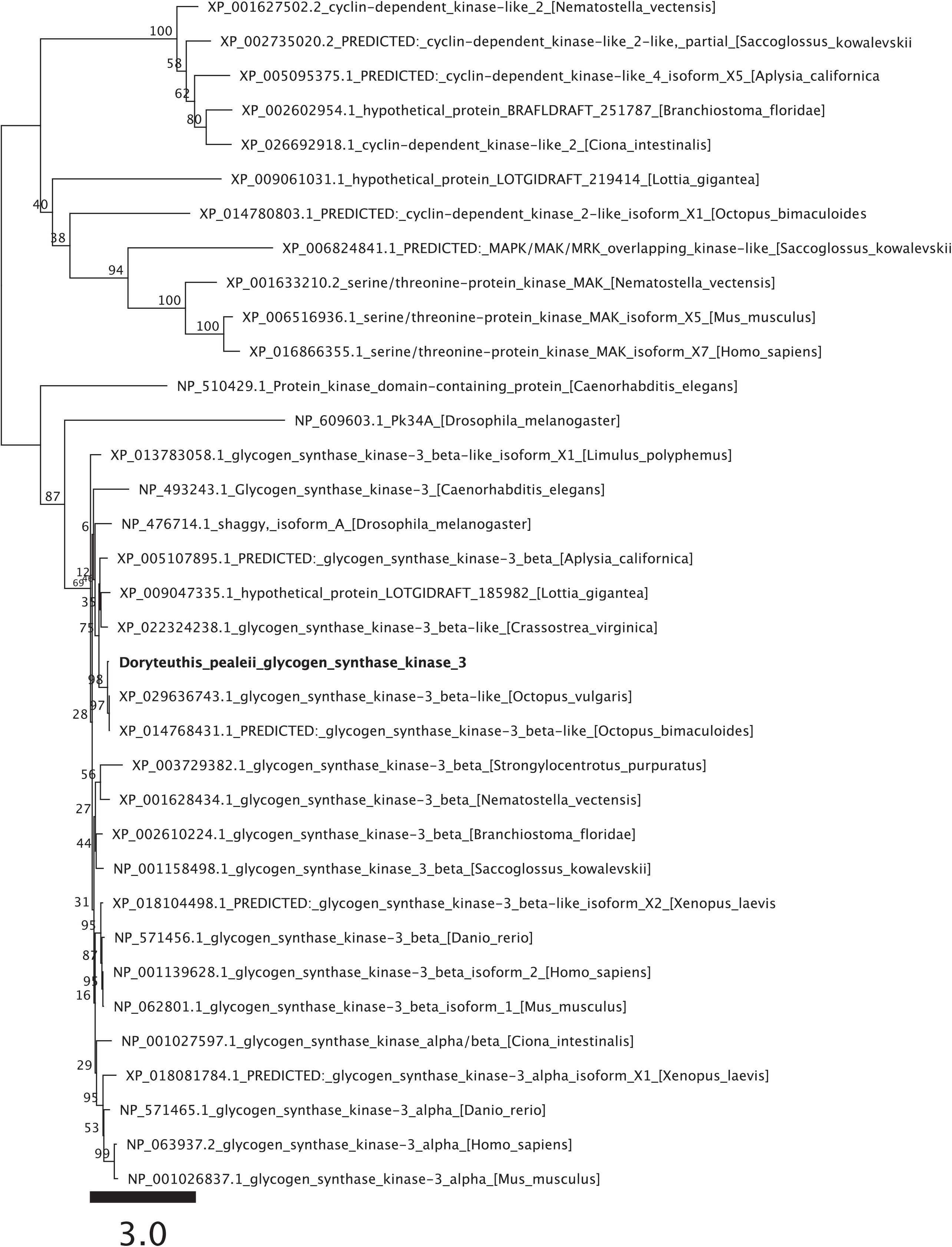
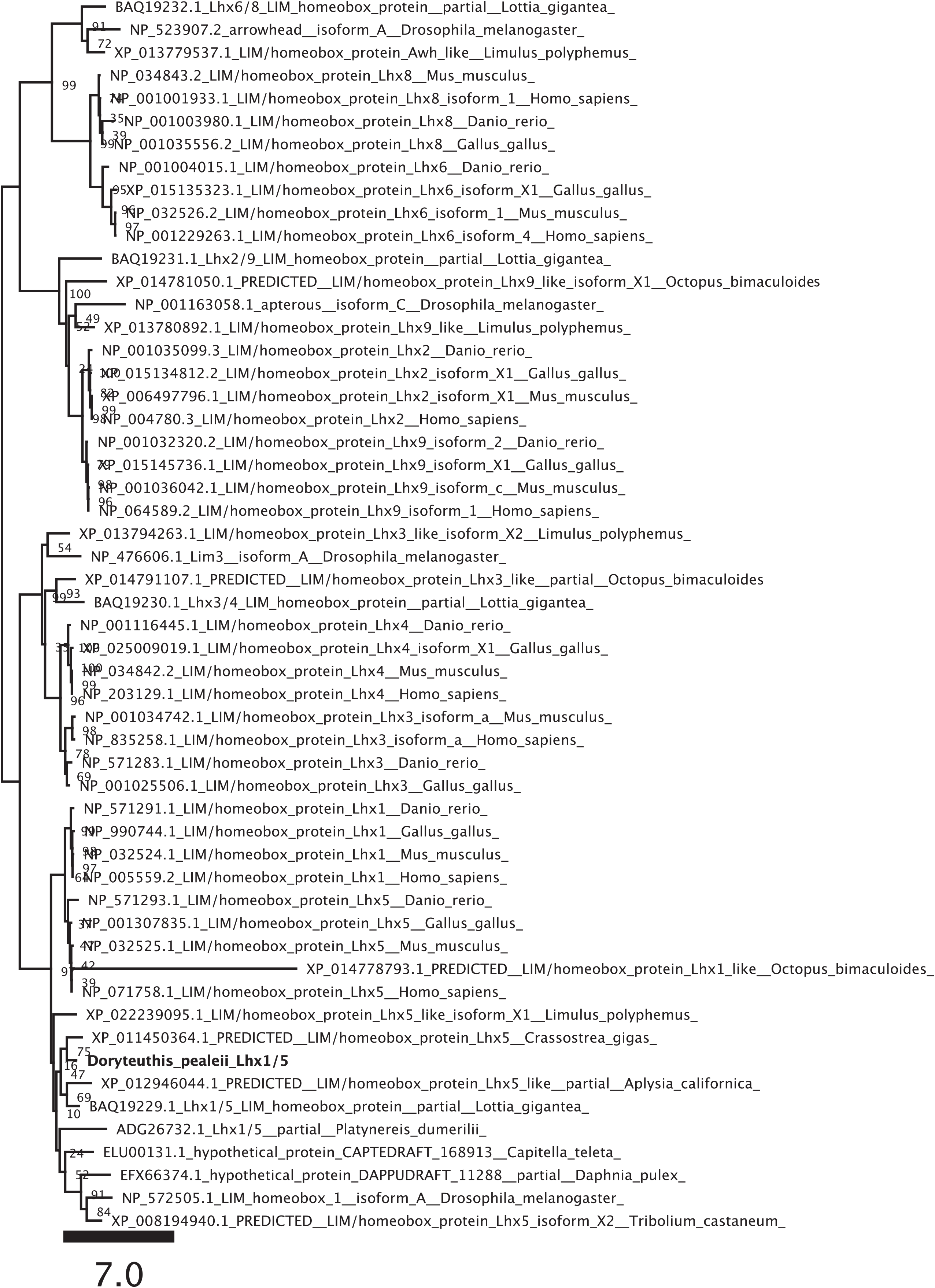
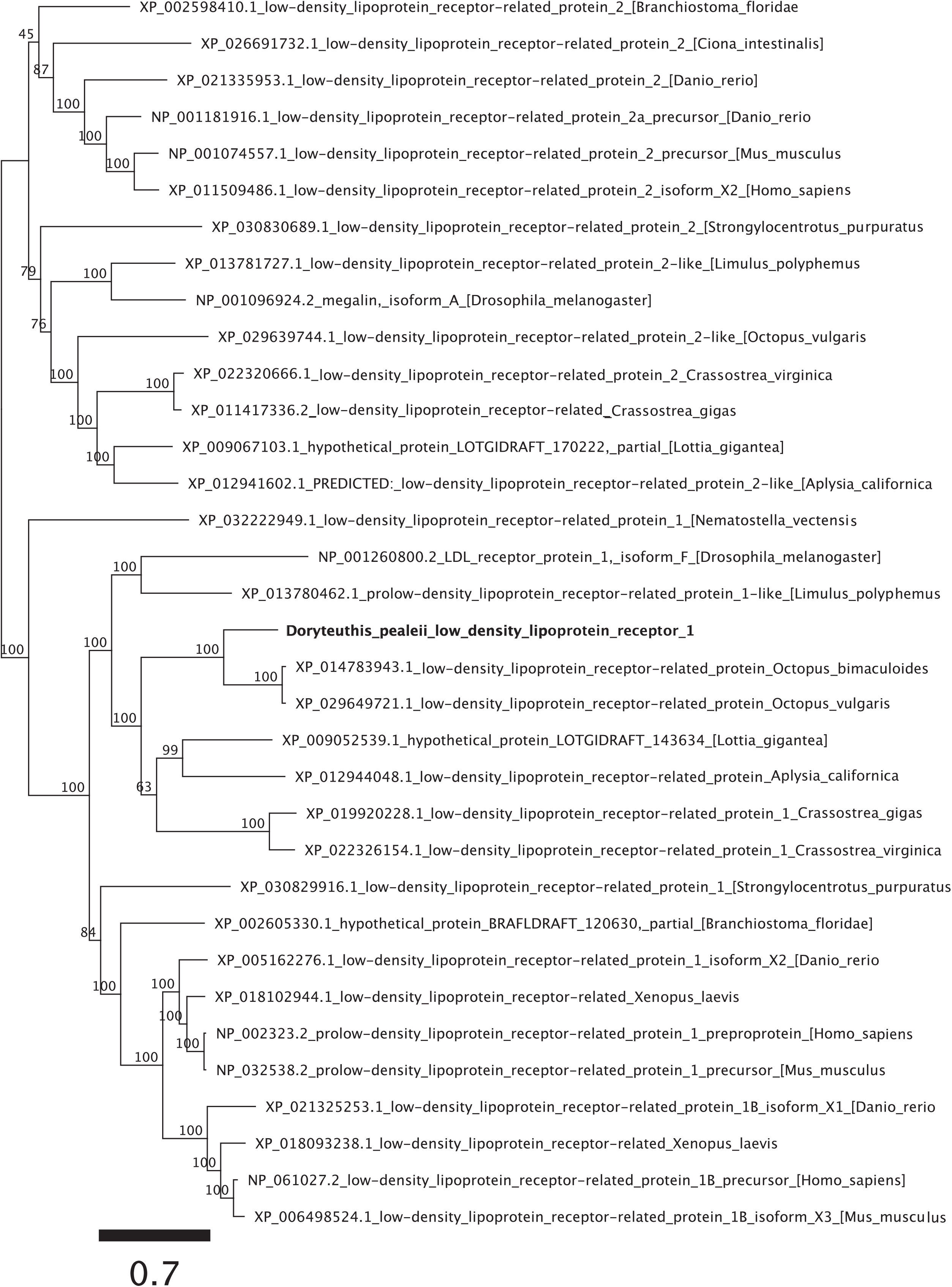
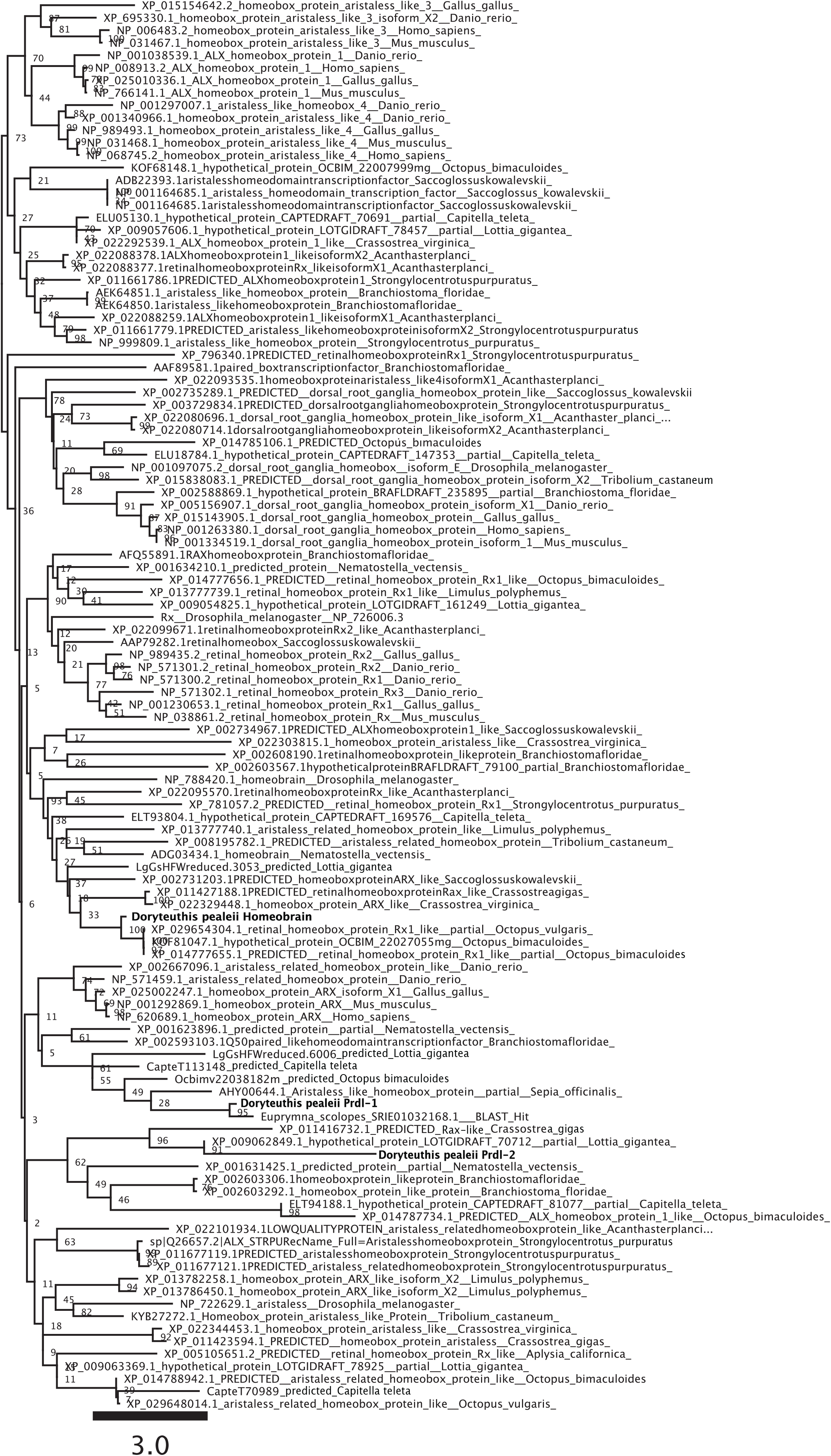
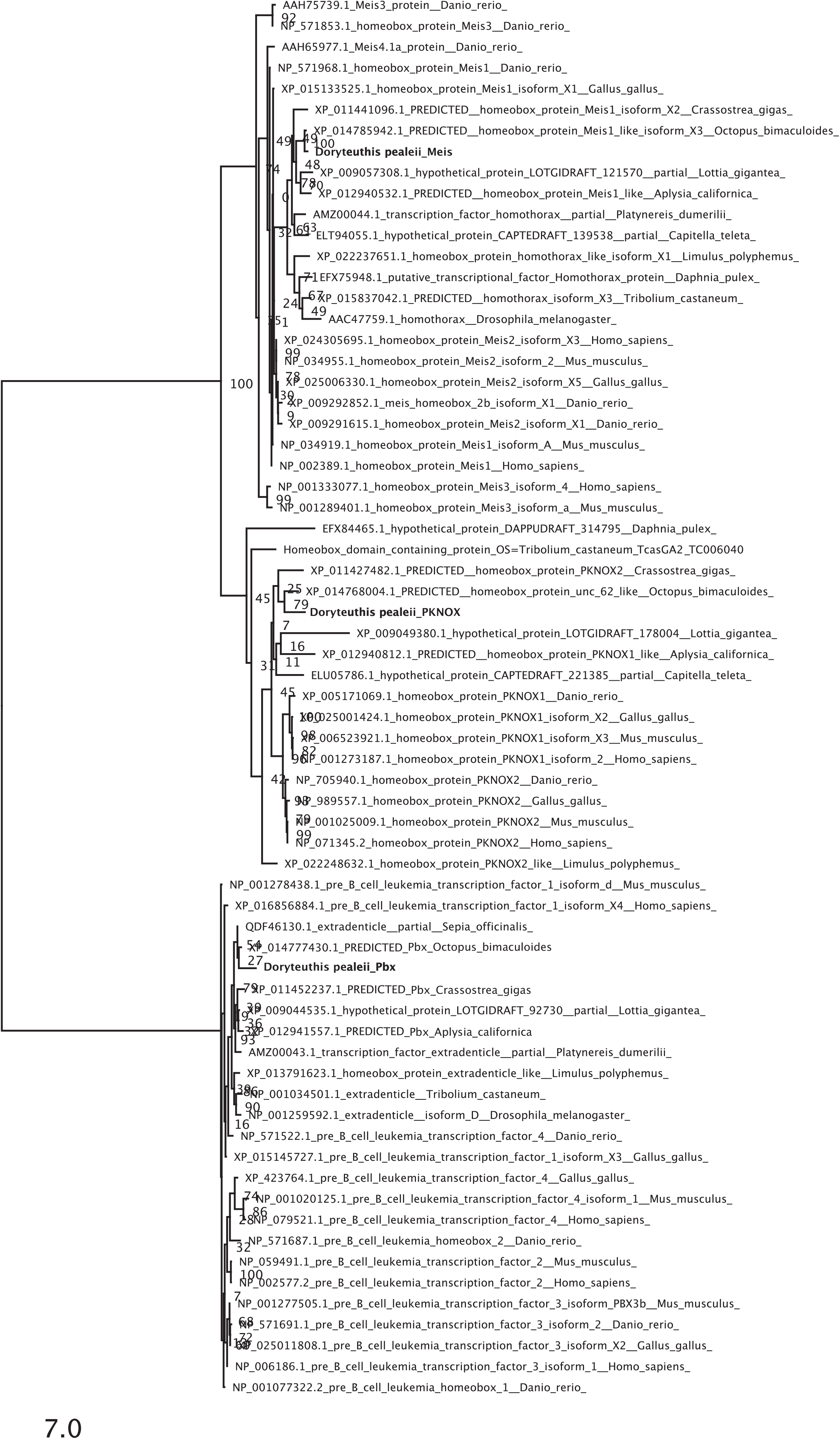
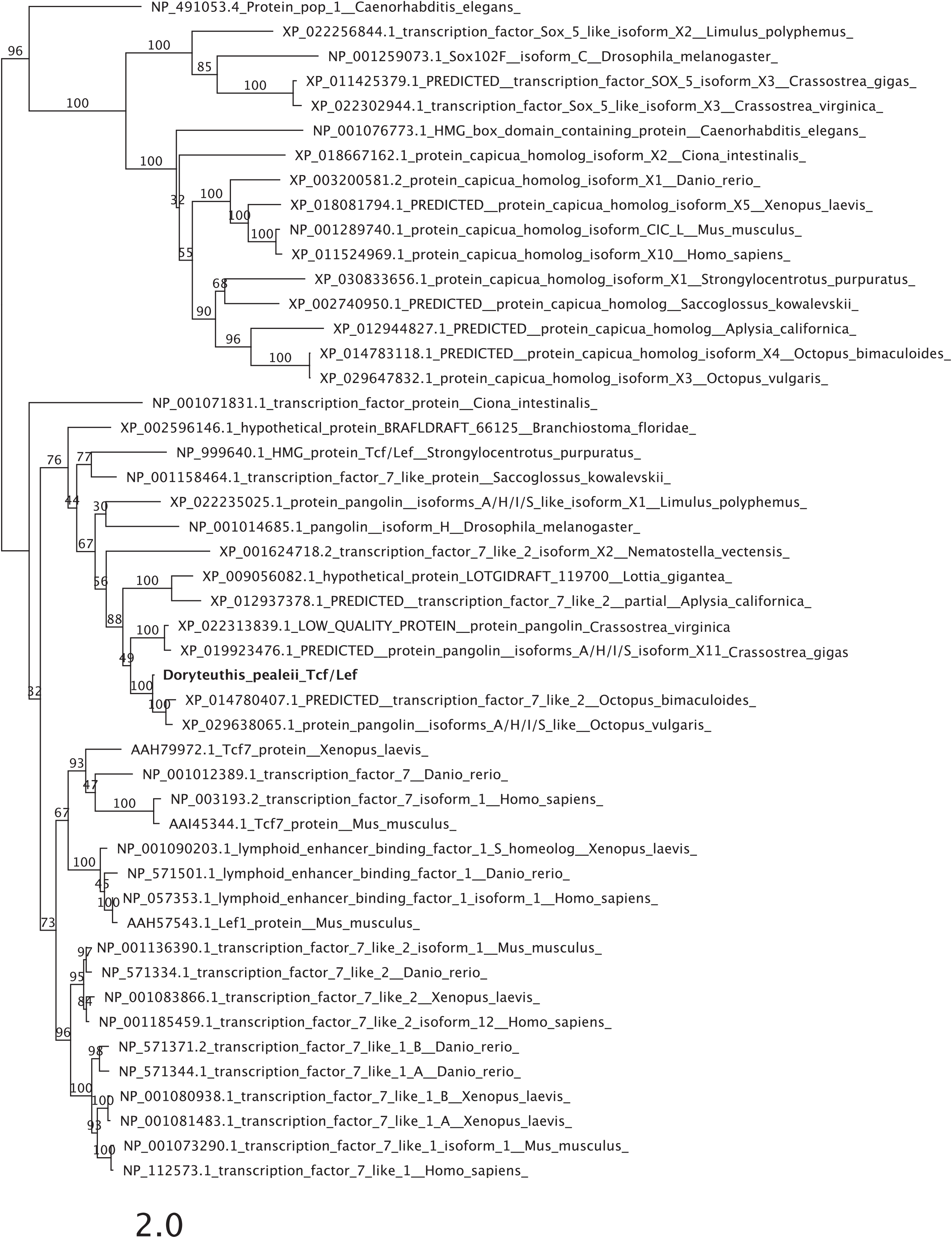
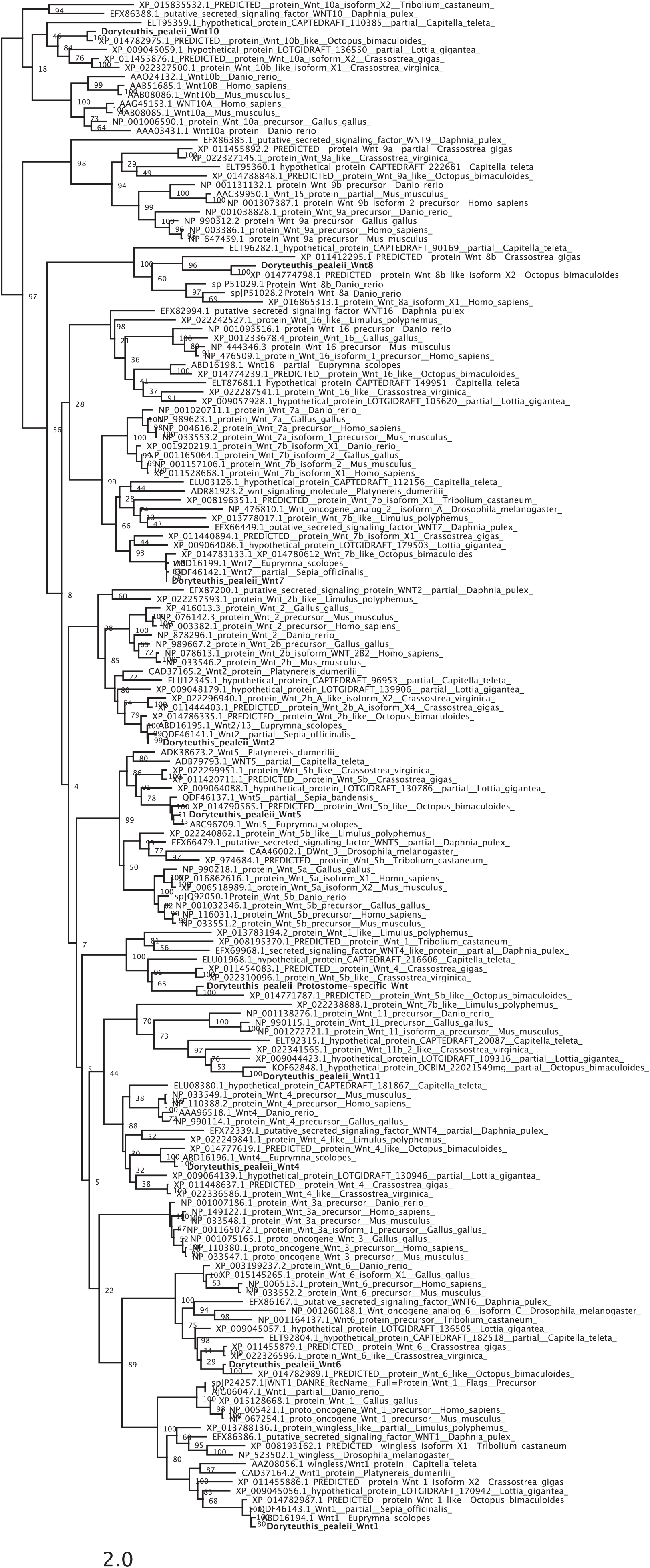
Maximum-likelihood phylogenetic trees for genes identified in this study.

**Sup Figure 2:**
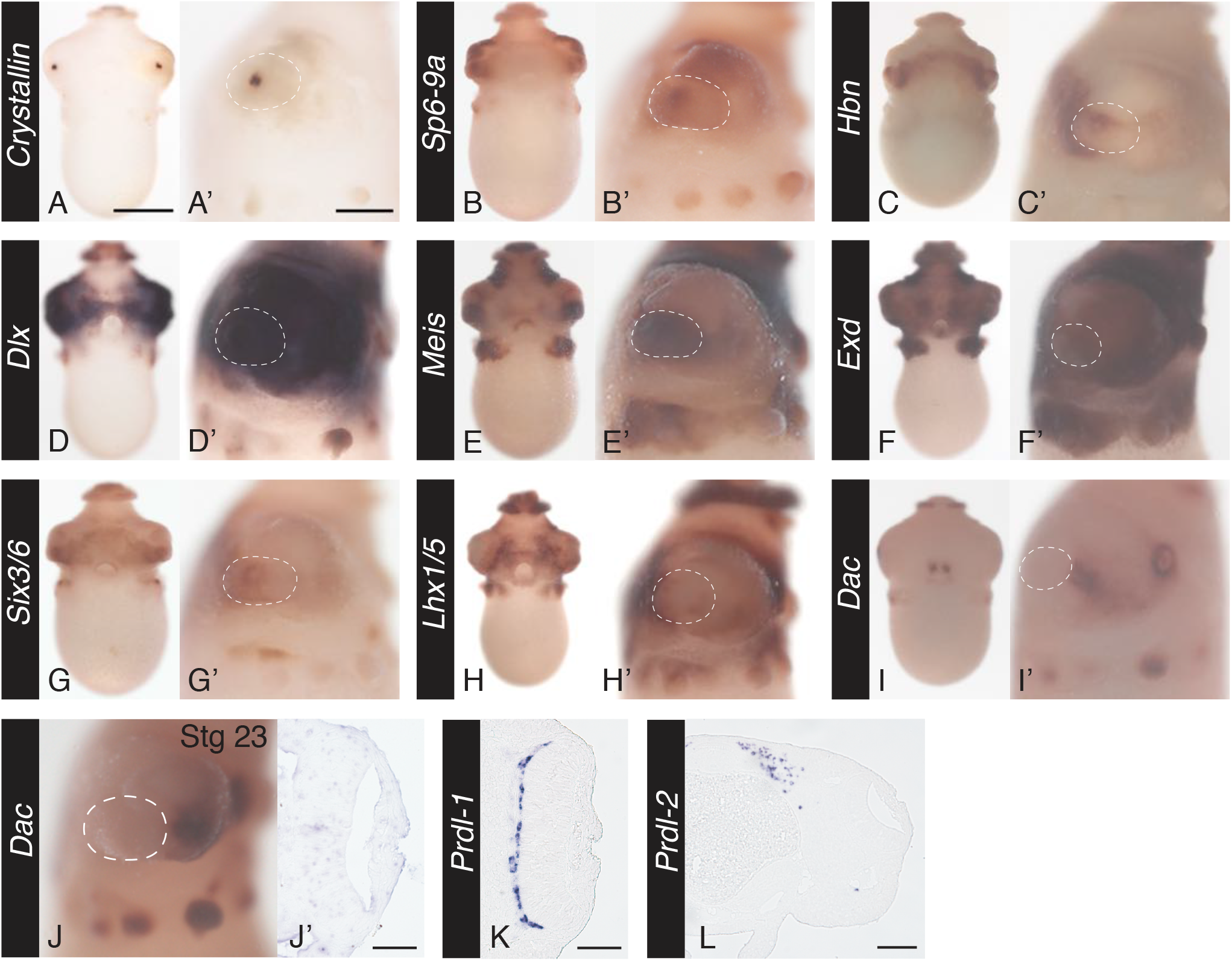
Limb network supplemental data. A-I) Gene expression at stage 21 for limb network genes. For all genes from left to right, Anterior whole-mount and lateral whole-mount, anterior to the left. Scale for whole-mount anterior view is 500 microns. Scale for lateral whole-mount view 200 microns. J, J’) Stage 23 Dac expression. J) Lateral whole mount, anterior to the left. J’) Sectioned image of the eye. Anterior is down. K & L) Sectioned image of expression of Prdl-1and Prdl-2. Scale is 50 microns on eye sections, 100 microns on brain section (Prdl-2)

**Sup Figure 3:**
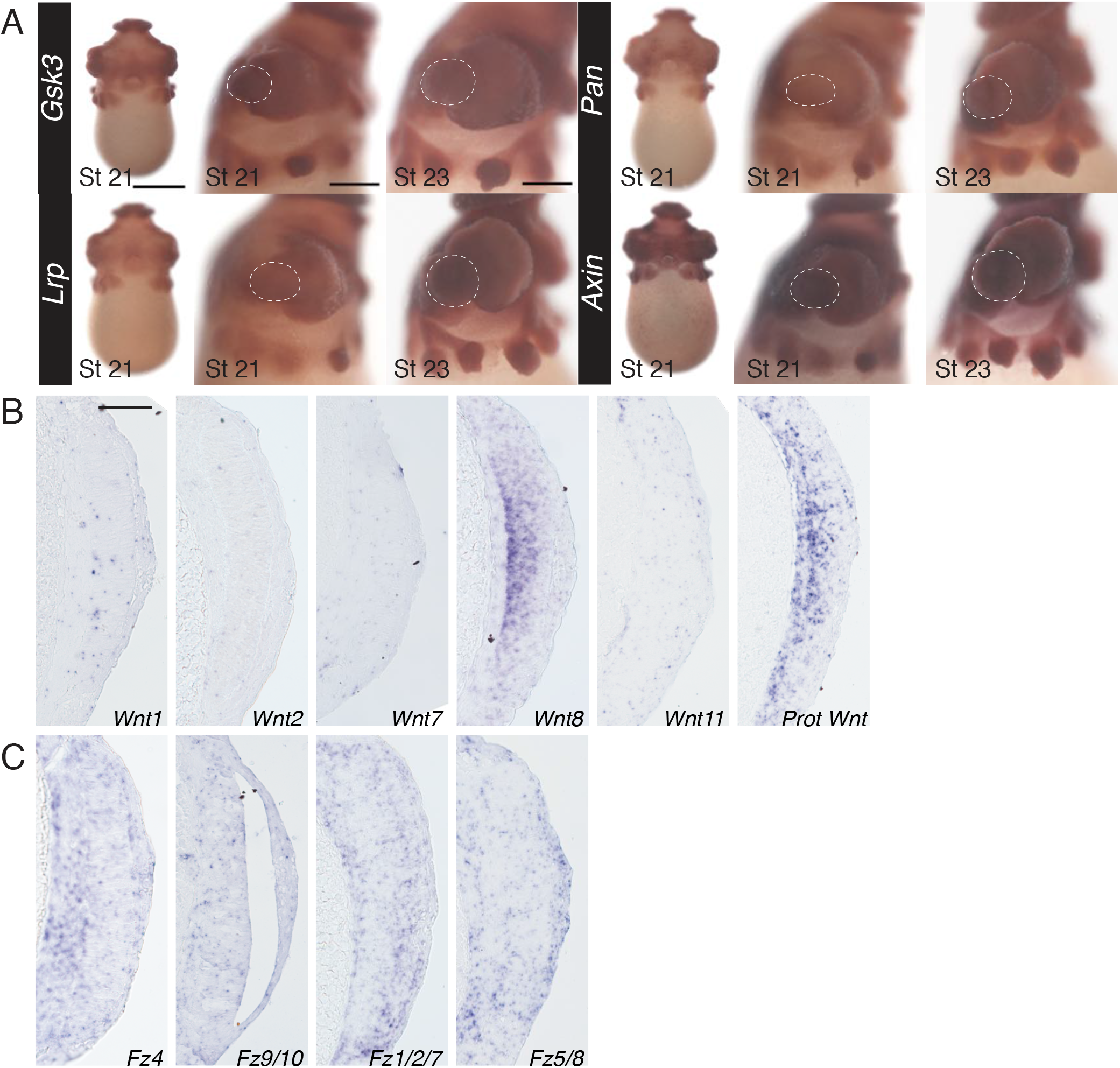
Wnt signaling expression supplemental data. A) Wnt signaling pathway member expression, *Gsk3, Lrp, Pan*, and *Axin*, at stage 21 and 23 in whole-mount. Anterior view of stage 21and lateral views of stage 21 and stage 23 (anterior to the left). B) Wnt gene expression at stage 21 in section. Anterior is down. C) Fz receptor gene expression at stage 21. Anterior is down. Scale for whole-mount anterior view is 500 microns. Scale for lateral whole-mount view 200 microns. Scale for sectioned images 50 microns.

**Sup Figure 4:**
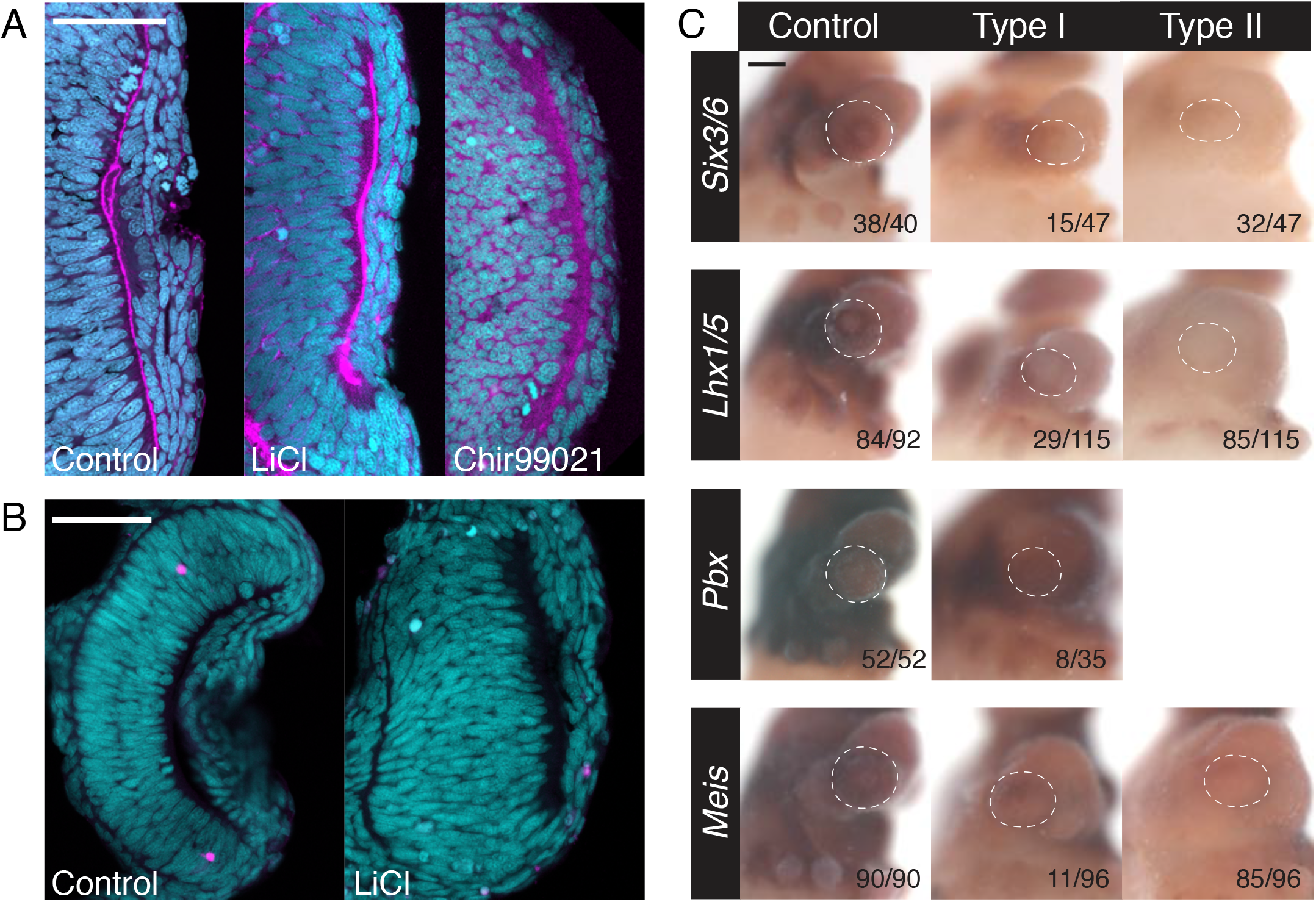
Wnt agonist experiment supplemental data. A) Wnt agonist experiments starting at stage 21. Embryos were treated for 24 hours and fixed immediately. LiCl and Chir99021 show similar phenotypes: Lack of anterior segment thickness and loss of lens formation. Sytox nuclear stain in cyan, Phalloidin stain in magenta. Scale is 50 microns. B) Tunel staining of the eye of Control and LiCl treated embryos. Sytox nuclear stain in cyan, Tunel stain in magenta. Similar amounts of cell death observed in control and treated animals. Scale is 100 microns C) *In situ* hybridization of limb patterning program members and and anterior segment markers after LiCl treatment. Type I (mild) and Type II (severe) phenotype. White dotted line outlines the eye in the lateral image. Number of eyes scored in control and the two phenotypes found in LiCl treated animals in the bottom right corner. Scale for lateral whole-mount view 200 microns.

## Notes

### Competing Interest Statement

The authors have declared no competing interest.

